# Global analysis of protein and small-molecule substrates of ubiquitin-like proteins (UBLs)

**DOI:** 10.1101/2024.12.05.626486

**Authors:** Guang-Can Shao, Zhen-Lin Chen, Shan Lu, Qing-Cui Wu, Yao Sheng, Jing Wang, Yan Ma, Jian-Hua Sui, Hao Chi, Xiang-Bing Qi, Si-Min He, Li-Lin Du, Meng-Qiu Dong

## Abstract

**Abstract:** Ubiquitin-like proteins (UBLs) constitute a family of evolutionarily conserved proteins that share similarities with ubiquitin in 3D structures and modification mechanisms. For most UBLs including Small-Ubiquitin-like Modifiers (SUMO), their modification sites on substrate proteins cannot be identified using the mass spectrometry-based method that has been successful for identifying ubiquitination sites, unless a UBL protein is mutated accordingly. To identify UBL modification sites without having to mutate UBL, we have developed a dedicated search engine pLink-UBL on the basis of pLink, a software tool for identification of cross-linked peptide pairs. pLink-UBL exhibited superior precision, sensitivity, and speed than “make-do” search engines such as MaxQuant, pFind, and pLink. For example, compared to MaxQuant, pLink-UBL increased the number of identified SUMOylation sites by 50 ∼ 300% from the same datasets. Additionally, we present a method for identifying small-molecule modifications of UBLs. This method involves antibody enrichment of a UBL C-terminal peptide following enrichment of a UBL protein, followed by LC-MS/MS analysis and a pFind 3 blind search to identify unexpected modifications. Using this method, we have discovered non-protein substrates of SUMO, of which spermidine is the major one for fission yeast SUMO Pmt3. Spermidine can be conjugated to the C-terminal carboxylate group of Pmt3 through its N^1^ or also likely, N^8^ amino group in the presence of SUMO E1, E2, and ATP. Pmt3-spermidine conjugation does not require E3 and can be reversed by SUMO isopeptidase Ulp1. SUMO-spermidine conjugation is present in mice and humans. Also, spermidine can be conjugated to ubiquitin *in vitro* by E1 and E2 in the presence of ATP. The above observations suggest that spermidine may be a common small molecule substrate of SUMO and possibly ubiquitin across eukaryotic species.

**Highlights:**

1. A specialized software pLink-UBL enables precise identification of UBL modification sites on protein substrates.
2. pFind 3 blind search enables identification of unexpected small-molecule substrates of a UBL protein.
3. Spermidine is a small molecule substrate of fission yeast SUMO Pmt3 as well as mammalian SUMO proteins.
4. The C-terminal carboxyl group of Pmt3 can be attached to the N^1^ or likely also the N^8^ amino group of spermidine in the presence of E1, E2, and ATP, and can be detached by SUMO isopeptidase.

## Introduction

Ubiquitin-like proteins (UBLs) are a family of evolutionarily conserved proteins that share with ubiquitin a characteristic β-grasp fold, also known as the ubiquitin super fold [1–4]. UBLs use a modification mechanism similar to that of ubiquitin, which involves a cascade of E1 (Ub/UBL-activating enzyme) [5], E2 (Ub/UBL-conjugating enzyme) [6, 7], and E3 (Ub/UBL ligase enzyme) [8] enzymes to catalyze the formation of an isopeptide bond between the carboxylate terminus of Ub or UBL and the side-chain [-amino group of a lysine residue in an acceptor protein, also called a substrate protein [9–11].

Since the discovery of the first UBL gene in 1979 [12], over 10 UBLs have been identified across plant and animal kingdoms [13–17], including Small Ubiquitin Modifiers (SUMO), Nedd8, Urm1, and Atg12. UBL modifications are implicated in diverse biological processes such as DNA damage repair, inflammation, autophagy, and oxidative stress response [18–24]. However, identification of UBL modification sites remains a challenge. This is because after trypsin digestion, which is a standard sample preparation step for mass spectrometry (MS)-based proteomics analysis, most UBLs leave a long C-terminal peptide (**Fig 1A**, 30 amino acids in the case of human SUMO2) covalently attached to what is left of a substrate protein—a peptide with a UBL-modified lysine residue and referred to in this study as a substrate peptide. A long remnant peptide of UBL complicates the fragmentation spectrum of a substrate peptide and thus interferes with the identification of the substrates peptide by conventional proteomics search engines [25–27]. Of the search engines in this category, MaxQuant [25] is used more often than others. MaxQuant allows users to define a PTM mass, such as that of a UBL C-terminal peptide, and to also define a number of *b* ions that could be generated from the UBL C-terminal peptide as signature ions [28–33]. However, there is a limit to the effectiveness of this strategy. Hence, mutagenesis is often used to generate a shorter UBL C-terminal peptide after trypsin digestion, sometimes just a diglycine peptide, in order to make use of the sophisticated tool kit for ubiquitination analysis. Although converting a long remnant peptide of UBL to diglycine facilitates MS data analysis, it opens the door to falsely identifying ubiquitinated peptides as UBL-modified peptides.

**Fig. 1.**
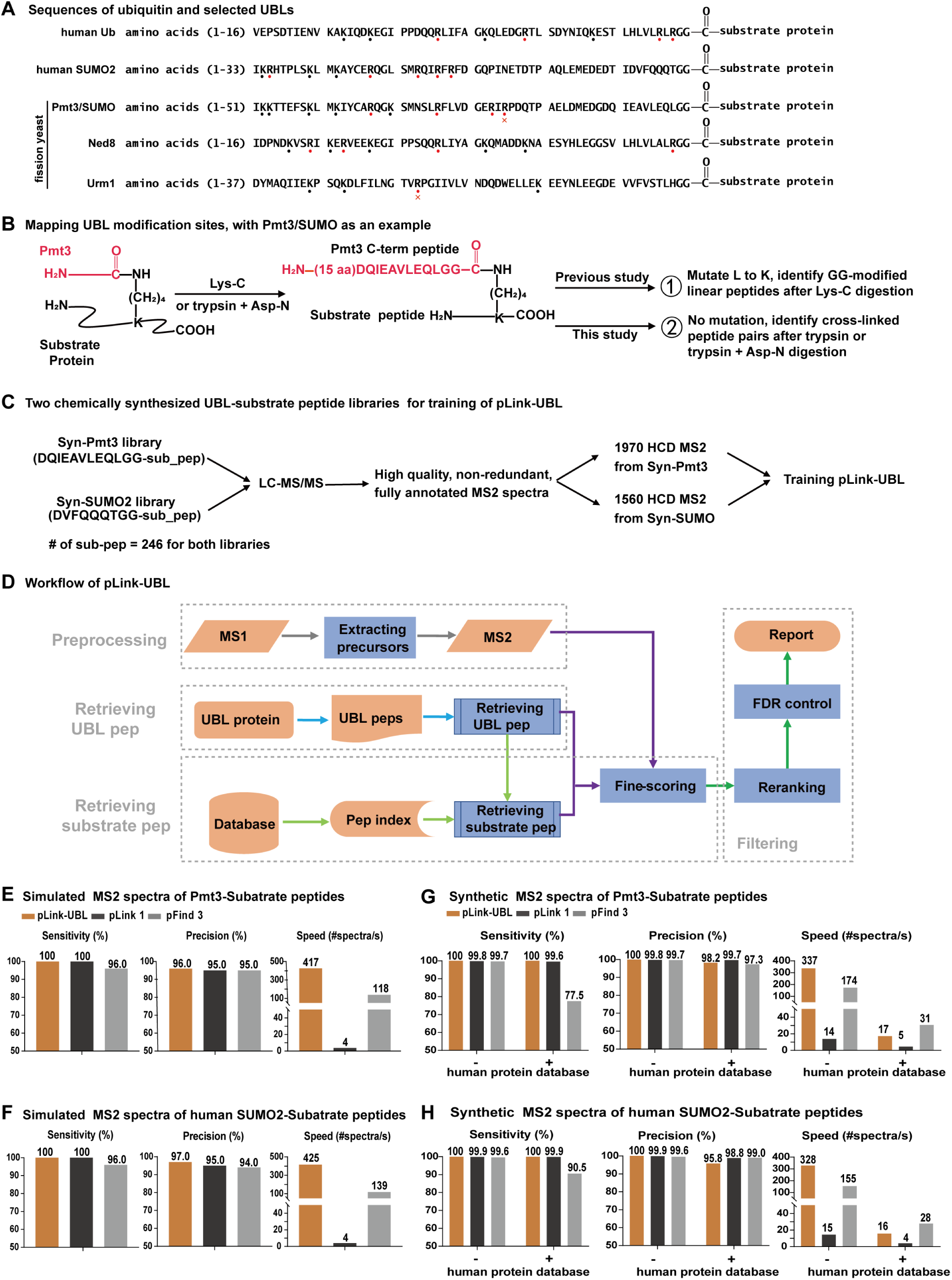
Development of pLink-UBL for identification of UBL modified proteins and sites. (A) Sequences of ubiquitin and selected UBLs, aligned from the C-terminus. **(B)** Strategies for proteomic profiling of UBL modification sites on substrate proteins, using fission yeast SUMO Pmt3 as an example. **(C)** Two synthetic UBL-substrate peptide libraries and the resulting training data for pLink-UBL. For the syn-Pmt3 library, the UBL sequence corresponds to the Pmt3 C-terminal peptide after trypsin+Asn-N digestion. For the syn-SUMO2 library, the UBL sequence corresponds to the human SUMO2/3 C-terminal peptide after trypsin+Asn-N digestion. **(D)** Workflow of pLink-UBL**. (E and F)** Performance evaluation of pLink-UBL on simulated MS2 of Pmt3-substrate peptides, and SUMO2-substrate peptides, respectively. **(G and H)** Performance evaluation of pLink-UBL on experimental MS2 generated from (C), the synthetic peptide libraries syn-Pmt3 and syn-SUMO2, respectively.

Rarely, a UBL such as Nedd8 leaves on the substrate peptide only a di-glycine remnant **(Fig 1A)**, which can be treated as a small post-translational modification (PTM) that adds114.0429 Da to the modified lysine residue in the substrate peptide. Such peptides can be readily identified by conventional proteomics search engines [34]. However, the downside is that ubiquitination also leaves a di-glycine remnant on substrate peptide after trypsin digestion [35, 36]. So, there is a concern of ambiguity—if Neddylated protein samples are contaminated with ubiquitinated proteins, di-glycine modified peptides could originate from either neddylation or ubiquitination. Such mistakes could be avoided if substrate peptides modified by a remnant UBL peptide are treated as cross-linked peptide pairs, and identified using cross-link search engines [37, 38]. In the case of Nedd8, a protease which is not trypsin may be used to generate a remnant C-terminal peptide longer than di-glycine. This approach has been explored in the past by ChopNspice [39]. ChopNspice converts the cross-linked peptide pairs to linear peptides, by concatenating the sequence of a remnant UBL peptide to the N-terminus of a substrate peptide. This is effective to some extent, but sensitivity and precision are compromised because a subset of experimental fragment ions are ignored and a subset of theoretical fragment ions are false.

The carboxylate termini of UBLs can be conjugated covalently to small molecule metabolites [40, 41]. For example, autophagy-related protein 8 (ATG8), a UBL in yeast cells, can be conjugated to phosphatidylethanolamine (PE) through the action of (E1) ATG7, (E2) ATG3 and (E3) the ATG12-ATG5-ATG16 complex [23, 42–44]. Likewise, conjugation of LC3—(homolog of ATG8) to PE is crucial for autophagosome formation in mammalian cells [45, 46].

In yeast cells, ubiquitin also modifies PE on endosomes and vacuoles (yeast lysosome) [47]. This is mediated by Uba 1 (E1), Ubc4/Ubc5 (E2), and Tul 1 (E3), and facilitates the recruitment of ESCRT components. Additionally, ubiquitin can modify lipopolysaccharides (LPS) on infecting *Salmonella* cells [48], which helps eliminate invading the bacteria from host cells. The hydroxyl or phosphate group of the lipid A moiety of LPS is suggested as a possible site of ubiquitination.

In this study, we aim to develop methods for analysis of proteins and non-protein substrates of UBLs. For the former, we have developed pLink-UBL based on the framework of pLink [37, 38], a search engine for identification of cross-linked peptide pairs. pLink-UBL treats UBL-modified substrate peptides as a special case of cross-linked peptide pairs, in which the sequence of one peptide (the substrate peptide) is variable and the other is not (the C-terminal peptide of a given UBL). To identify non-protein substrates of UBLs in an unbiased way, we resorted to the blind search function of pFind 3 [49, 50] and looked for unexpected, high-frequency modifications on a UBL C-terminal peptide. To prove the concept, we developed an antibody to enrich the C-terminal peptide of Pmt3, the fission yeast SUMO. MS analysis of the peptides enriched by this antibody and pFind 3 blind search led to the discovery that the C-terminus of Pmt3 is often conjugated to spermidine, and likely to several other polyamines at a lower frequency. We further demonstrated that Pmt3 is conjugated to the N^1^ or also likely, N^8^ amino group of spermidine, and this can be catalyzed by E1 and E2 in the presence of ATP. Conversely, Pmt3 can be detached from spermidine by Ulp1, the fission yeast SUMO isopeptidase [51, 52]. From public MS data depositories, SUMO-spermidine conjugation also occurs in mice and humans.

### Experimental Procedures Reagents

Lys-C (cat. #VA1170), Trypsin (cat. #V5280), and Asp-N (cat. #V1621) were purchased from Promega. The anti-HA antibody was purchased from MBL (cat. #M180-3), the anti-GST antibody from Abmart (cat. #AbM59001), and the mouse anti-ubiquitin antibody from ThermoFisher (cat. #14-6078-82) and the HRP-conjugated goat anti-mouse IgG from Sigma Aldrich (cat. #A4416). The anti-Pmt3 rabbit polyclonal antibody was produced at the NIBS antibody facility using recombinant Pmt3 as the antigen. The Ni^2+^-NTA agarose beads were purchased from QIAGEN (cat. #30210). DMP (dimethyl pimelimidate) was purchased from ThermoFisher (cat. #21667). Protein A agarose beads were purchased from Smart-Lifesciences (cat. #SA023005).

### Fission yeast strains, plasmids, and growth conditions

Fission yeast strains and plasmids used in this study are detailed in **Supplementary Table S1 and Table S2**. Fission yeast strains DY8577, SY509, SY535, and SY563 were cultured in liquid YES medium at 30°C and harvested by centrifugation when the optical density at 600 nm (OD600) reached approximately 1.5. The temperature-sensitive *slx8-1* mutant was grown initially at 30°C and subsequently shifted to 37°C for 12 hours before harvest. The harvested cells—in pellet forms—were washed twice with ice-cold PBS and subsequently flash-frozen using liquid nitrogen.

### Cell lysis, protein extraction, and nickel affinity purification

A total of 1500 OD fission yeast cells were lysed using a TissueLyser-24 and subsequently resuspended in 15 mL of lysis buffer (6 M guanidine-HCl; 0.1 M PBS, pH 8.0; 0.01 M Tris-HCl, pH 8.0; 10 mM imidazole, pH 8.0). The proteins were allowed to be solubilized in lysis buffer on a roller mixer at 4°C for 1 h. After centrifugation at 12000 rpm for 30 min, the clarified lysate was mixed with 200 µL of Ni^2+^-NTA agarose beads and incubated at 4°C for 4 h. Subsequently, the beads were collected and washed with 1 mL of each buffer in the following order: lysis buffer for three times, wash buffer pH 8.0 (8 M urea; 0.1 M PBS, pH 8.0; 10 mM Tris-HCl, pH 8.0; 15 mM imidazole) for three times, wash buffer pH 6.3 (8 M urea; 0.1 M PBS, pH 6.3; 10 mM Tris-HCl, pH 6.3) twice, and once more with wash buffer pH 8.0. Proteins were eluted in three sequential steps with 3ml of elution buffer (8 M urea; 0.1 M PBS, pH 8.0; 10 mM Tris-HCl, pH 8.0; 500 mM imidazole). The three elution fractions were combined and concentrated to approximately 100 µL using 30-kDa cut-off filter units.

For immunoblot analysis, 5 µL of protein sample was separated by SDS–polyacrylamide gel electrophoresis on a 12% gel followed by transfer to a PVDF membrane. After blocking with 5% non-fat milk, the membrane was incubated with a mouse anti-HA antibody (diluted 1:5,000) at 4°C for 1 h followed by a horseradish peroxidase-conjugated goat anti-mouse IgG secondary antibody (dilution: 1:5000) at room temperature for 1 h. The membrane was washed three times with TBST for 10 min each time. The SUMOylated species were detected using an ECL Plus kit (GE Healthcare).

### Development of antibody against the C-terminal peptide of Pmt3 (anti-IG27 antibody)

To generate an antibody that specifically recognizes the C-terminal peptide of Pmt3, a 27-aa peptide of Pmt3 (IG27) was selected for antigen preparation. This peptide sequence corresponds to the most C-terminal peptide of Pmt3 after trypsin digestion. To mimic the consensus motif of SUMOylation sites (ΨKXE, where Ψ denotes a hydrophobic amino acid, such as I, L, and V), the antigen peptide was designed as a conjugated peptide (V{K(IRPDQTPAELDMEDGDQIEAVLEQLGG)}AE). This antigen peptide was subsequently conjugated with keyhole limpet hemocyanin protein via the cysteine residue as an immunogen. Two rabbits were immunized using standard immunization of three injections. Ultimately, 40 mL of immune serum was obtained from one of the rabbits.

### Evaluate anti-IG27 antibody using IG27-GST

To evaluate the binding affinity of the immune serum, the fusion protein IG27-GST was expressed and purified in *E. coli* BL21. 15 µL of anti-IG27 immune serum was diluted 20-fold with PBS, and subsequently incubated with 2 µg of IG27-GST at 4°C for 1 h. Following this incubation, 10 µL of protein A agarose slurry was added to the mixture, which was further incubated at 4°C for an additional 3 h. Thereafter, the saturated protein A agarose beads were washed three times using 1×PBS. The IG27-GST was eluted using 50 µL of HU buffer (8 M urea; 200 mM Tris-HCl, pH 6.8; 1 mM EDTA; 5% (w/v) SDS; 0.1% (w/v) bromophenol blue; 1.5% (w/v) dithiothreitol). The eluted proteins were then separated through SDS–polyacrylamide gel electrophoresis on a 12% gel. Afterward, proteins were transferred to a PVDF membrane and subjected to immunoblotting using a mouse anti-GST antibody (1:5000).

### Antibody cross-linking

Anti-IG27 immune serum was diluted 20-fold with PBS and subsequently incubated with protein A agarose beads (15 µL of immune serum per 10 µL of protein A agarose slurry) at 4°C for 1 h. The saturated protein A agarose beads were then washed three times with PBS and 200 mM triethanolamine pH 8.3, respectively. For crosslinking, 10 µL of 5 mM DMP in 200 mM triethanolamine pH 8.3 was added to 1 µL of antibody-bound bead slurry and incubated at room temperature for 1 h. The reaction was quenched for 30 min using 50 mM Tris-HCl pH 8.0.

Cross-linked beads were washed three times with ice-cold PBS and stored in PBS containing 50% glycerol at -20°C for future use.

### Evaluate anti-IG27 antibody using synthetic SUMOylated peptides

To assess the efficacy of the anti-IG27 antibody in enriching synthetic SUMOylated peptides, we spiked about 7.5 µg of synthetic SUMOylated peptides into a complex background comprising about 72 µg of trypsin digests of total protein from fission yeast. Subsequently, the complex sample was subjected to a 3 h incubation with the anti-IG27 antibody at 4°C. The anti-IG27 antibody-bound beads were washed three times with 1 mL of 1×PBS, followed by two washes with 1 mL of 0.1×PBS. The SUMOylated peptides were eluted using 100 µL of 0.2% formic acid in water, which was repeated three times and combined. The eluted peptides were subsequently dried by speed vacuum and stored at -80°C for further MS spectrometry analysis. The mass spectrometry data were analyzed by the software pLink-UBL and pFind 3.

### Enrichment of SUMOylated peptides from fission yeast sample

To enrich SUMOylated peptides from the fission yeast sample, on-bead trypsin digestion was carried out following Ni^2+^-NTA agarose pull-down from 1500 OD cells. The ratio of trypsin to protein was 1:50 by weight, and this was done overnight at 37°C. The digested peptides were desalted using Pierce™ Peptide Desalting Spin Columns and dried by speed vacuum. The dried peptides were reconstituted in 500 µL of TBS containing 0.1% NP40. After centrifugation, 100 µL of antibody-bound bead slurry was added and incubated at 4°C for 1 h. The antibody-bound beads were washed three times with 1 mL of 1×PBS, followed by two washes with 1 mL of 0.1×PBS. The SUMOylated peptides were eluted three times with 100 µL of 0.2% formic acid in water and then combined. Finally, the eluted peptides were dried down by speed vacuum and stored at -80°C for further MS analysis.

### LC-MS/MS analysis

The digested peptides were analyzed using an EASY-nLC 1000 system interfaced with QE-HF mass spectrometer. Peptides were loaded onto a pre-column (100 μm ID, 5 cm long, packed with ODS-AQ 12 nm–10 μm beads) and separated on an analytical column (75 μm ID, 15 cm long, packed with Luna C18 1.9 μm 100 Å resin). Samples were separated with a 120 min linear reverse-phase gradient at a flow rate of 300 nL/min as follows: 10%-30% B in 95 min, 30%-80% B in 7 min, and 80% B for 16 min (A = 0.1% FA, B = 100% ACN, 0.1% FA). Spectra were acquired in data-dependent mode: the top 15 most intense precursor ions from each full scan (resolution 60,000) were isolated for HCD MS2 (resolution 30,000, HCD Collision Energy 30%) with a dynamic exclusion time of 30 s, a target AGC of 1E5 and a maximum injection time of 150 ms. Precursors with 1+, 2+, more than 7+, or unassigned charge states were excluded.

### Data Processing

The open search function of pFind 3 (3.2.0) was used for peptide identification. The number of maximum missed cleavage sites was set to 3. The tolerances of the precursor and the fragment were both set to 20 ppm. The false discovery rate was 1% at the peptide level.

The restrict search function of pFind 3 (3.2.0) was used for the identification of SUMO-spermidine conjugation or ubiquitin-spermidine conjugation. Carbamidomethyl(C) was set as a fixed modification, and Oxidation(M), Deamination(N), Carbamyl(Any N-term), and spermidine (G) were set as variable modifications. The number of maximum missed cleavage sites was set to 3. The tolerances of the precursor and the fragment were both set to 20 ppm. The false discovery rate was 1% at the peptide level.

The blind search function of pFind 3 (3.2.0) was used for the identification of novel non-protein substrates. The number of the maximum missed cleavage sites was set to 3. The tolerances of the precursor and the fragment were both set to 20 ppm. The false discovery rate was 1% at the peptide level.

MaxQuant (1.6.6.0), pFind 3 (3.0.11), and pLink 1 (1.23) were used for evaluation of pLink-UBL, Carbamidomethyl(C) was set as a fixed modification, and Oxidation(M), Deamination(N), and Carbamyl(Any N-term) were set as variable modifications. The number of maximum missed cleavage sites was set to 5. The tolerances of the precursor and the fragment were both set to 20 ppm. The false discovery rate was 5% at the spectra level.

### Recombinant protein production

6×His tagged recombinant proteins were expressed in *E. coli* BL21. The *E. coli* cells were cultured in 50 mL of LB medium (5 g/L yeast extract; 10 g/L NaCl; 10 g/L bacto tryptone) supplemented with 50 mg/L of ampicillin. When the optical density at 600 nm (OD_600_) reached 0.6–0.8, the expression of the recombinant proteins was induced by adding 1 mM IPTG at 16°C for 18 h. The cells were then harvested and resuspended in 8 mL of lysis buffer (50 mM NaH_2_PO_4_, pH 8.0; 300 mM NaCl; 10% glycerol; 10 mM imidazole). Subsequently, the cells were lysed by sonication in a 10 mL beaker, and the resulting lysate was centrifuged at 12000 rpm for 15 min at 4°C. The supernatant was incubated with Ni^2+^-NTA agarose beads at 4°C for 1 h. The beads were washed three times with wash buffer (50 mM NaH_2_PO_4_, pH 8.0; 300 mM NaCl; 10% glycerol; 20 mM imidazole). The proteins were eluted using 500 µL of elution buffer (50 mM NaH_2_PO_4_, pH 8.0; 250 mM NaCl; 10% glycerol; 250 mM imidazole). The eluted protein was stored at −80°C for future use.

For *in vitro* SUMO-spermidine conjugation, 6×His-GST-Rad31_Fub2 (Rad31 and Fub2 cDNA were fused together with a short linker), 6×His-GST-Hus5, 6×His-GST-Pli1, and 6×His-Pmt3 were purified from *E. coli* BL21 **(Supplementary Fig. S1A and S1B)** [53]. For evaluation of the anti-IG27 antibody, GST and IG27-GST were purified from *E. coli* BL21 **(Supplementary Fig. S1C)**.

For *in vitro* ubiquitin-spermidine conjugation, HA-Ubiquitin, 6×His-Uba1, 6×His-Ubc1, 6×His-Ubc4, 6×His-Ubc6, 6×His-Ubc7, 6×His-Ubc8, 6×His-Ubc11, 6×His-Ubc13, 6×His-Ubc14, 6×His-Ubc15, 6×His-Ubc16, 6×His-Mms2, 6×His-Rhp6 were purified from *E. coli* BL21 **(Supplementary Fig. S1D)**.

### *In vitro* SUMO-spermidine conjugation and ubiquitin-spermidine conjugation

For *in vitro* SUMO-spermidine conjugation, 10 mM spermidine was incubated with 2 µg of Rad31_Fub2, 2 µg of Hus5, 2 µg of Pli1, and 1 µg of Pmt3 in a reaction buffer (50 mM Tris-HCl, pH 7.5; 100 mM NaCl; 10 mM MgCl_2_; 5 mM ATP; 1 mM DTT; 0.2 mM CaCl_2_) at 30°C overnight in a total volume of 20 µL. The reaction was stopped by adding HU loading buffer and then separated by SDS–polyacrylamide gel electrophoresis after boiling for 5 min. To enhance protein binding to the PVDF membrane, the membrane was treated with 1% glutaraldehyde in PBS for 30 min immediately following the protein transfer step. After blocking with 5% non-fat milk, the membrane was incubated with the primary antibody (anti-Pmt3, dilution: 1:1500) followed by the secondary antibody (HRP-conjugated goat anti-rabbit IgG, dilution: 1:5000).

For *in vitro* ubiquitin-spermidine conjugation, 10 mM spermidine was incubated with 2 µg of Uba1, 2 µg of Ubc4 (or other E2s), and 1 µg of ubiquitin in 20 µL of reaction buffer (50 mM Tris-HCl, pH 7.5; 100 mM NaCl; 10 mM MgCl_2_; 5 mM ATP; 1 mM DTT; 0.2 mM CaCl_2_) at 30°C overnight. The reaction was stopped by adding HU loading buffer and then separated by SDS–PAGE after boiling for 5 min. To enhance protein binding to the PVDF membrane, the membrane was treated with 1% glutaraldehyde in PBS for 30 min immediately following the protein transfer step. After blocking with 5% non-fat milk, the membrane was incubated with the primary antibody (anti-ubiquitin, dilution: 1:1200) followed by the secondary antibody (HRP-conjugated goat anti-mouse IgG, dilution: 1:5000).

## Datasets

### Preparation of the simulated dataset and database search

Two simulated datasets were generated using pSimXL1 [37]: the Simulated Pmt3-substrate peptide dataset and the Simulated SUMO2-substrate peptide dataset. Each dataset consists of 10,000 spectra from the SUMOylation of 1,000 *E. coli* proteins. To generate higher-quality simulated spectra, both occurrence probabilities and intensity distributions of fragment ions were set as those analyzed from synthetic datasets **(Supplementary Fig. S2)**, making the simulated spectra as high-quality as synthetic spectra. **Supplementary Fig. S3** shows two examples of simulated Pmt3 and SUMO2-modified spectra. Both SUMO and substrate peptides are *in silico* generated only 1^+^ and 2^+^ *b/y* ions, the simulated spectra are rather simpler compared to real-world spectra, and all search engines were expected to achieve high sensitivity and precision. We thus used the simulated datasets as a qualification test.

Database searches were performed with trypsin specificity and three missed cleavage sites were allowed. Peptides were accepted with a length range between 6 and 60 amino acids and a mass range between 600 and 6,000 Da. Oxidation [M] was chosen as a variable modification. Mass tolerances for the precursor were set to 10 ppm, and for the fragment, they were set to 20 ppm to search against the first 1,000 proteins in the *E. coli* database. For speed evaluation, search engines used eight threads for the search, and the computing times were measured on the same computer equipped with (Windows Server, Intel Xeon E5-2670 CPU with 32 cores, 2.6 GHz,

128 GB RAM).

### Preparation of the synthetic dataset and database search

Two synthetic datasets were generated: Synthetic Pmt3-substrate peptide and Synthetic SUMO2-substrate peptide, which were chemically SUMOylated by Pmt3 and SUMO2, respectively, using 246 substrate peptides from 30 templates. Ultimately, we successfully obtained 492 SUMOylated peptides through a condensation reaction involving the carboxyl group of the C-terminal glycine of SUMO and the side-chain ɛ-amino group of lysine residues in the substrate peptides **(Supplementary Fig. S4)**. The Synthetic Pmt3-substrate peptide dataset consisted of fully annotated 1,970 positive PSMs and equivalent negative PSMs, while the Synthetic SUMO2-substrate peptide dataset consisted of fully annotated 1,560 positive PSMs and equivalent negative PSMs.

Database search was performed with trypsin specificity and three missed cleavage sites were allowed. Peptides were accepted with a length range between 6 and 60 amino acids and a mass range between 600 and 6,000 Da, Oxidation[M] was chosen as a variable modification. Mass tolerances for the precursor were set to 10 ppm, and for the fragment, they were set to 20 ppm to search against the synthetic peptide database (494 peptides) and the human database (92,952 proteins). For speed evaluation, search engines used eight threads for the search, and the computing times were measured on the same computer equipped with (Windows Server, Intel Xeon E5-2670 CPU with 32 cores, 2.6 GHz, 128 GB RAM).

### Preparation of the fission yeast dataset and database search

A total of 3000 OD fission yeast cells were lysed using a TissueLyser-24 and subsequently resuspended in 30 mL of lysis buffer (6 M guanidine-HCl; 0.1 M PBS, pH 8.0; 0.01 M Tris-HCl, pH 8.0; 10 mM imidazole, pH 8.0). The proteins were allowed to be solubilized in lysis buffer on a roller mixer at 4°C for 1 h. After centrifugation at 12000 rpm for 30 min, the clarified lysate was mixed with 400 µL of Ni^2+^-NTA agarose beads and incubated at 4°C for 4 h. Subsequently, the beads were collected and washed with 1 mL of each buffer in the following order: lysis buffer for three times, wash buffer pH 8.0 (8 M urea; 0.1 M PBS, pH 8.0; 10 mM Tris-HCl, pH 8.0; 15 mM imidazole) for three times, wash buffer pH 6.3 (8 M urea; 0.1 M PBS, pH 6.3; 10 mM Tris-HCl, pH 6.3) twice, and once more with wash buffer pH 8.0. Proteins were eluted in three sequential steps with 3ml of elution buffer (8 M urea; 0.1 M PBS, pH 8.0; 10 mM Tris-HCl, pH 8.0; 500 mM imidazole). The three elution fractions were combined and concentrated to approximately 100 µL using 30-kDa cut-off filter units.

The concentrated sample underwent overnight digestion at 37°C using a combination of trypsin and Asp-N enzymes before SCX fractionation. Seven SCX fractions were collected by sequential elution using different concentrations of ammonium acetate (25 mM, 50 mM, 75 mM, 100 mM, 250 mM, 500 mM, and 1 M).

The LC-MS/MS analysis was performed on a Q-Exactive HF mass spectrometer coupled to an Easy-nLC 1000 system. Peptides were loaded on a pre-column (100 μm ID, 5 cm long, packed with ODS-AQ 120 Å–10 μm beads from YMC Co., Ltd) and further separated on an analytical column (75 μm ID, 15 cm long, packed with C18 1.9 μm 100 Å resin from Welch Materials). Samples were separated with a 120 min linear reverse-phase gradient at a flow rate of 200 nL/min as follows: 0%-30% B in 100 min, 30-80% B in 7 min, and 80% B for 13 min (A =0.1% formic acid in H_2_O, B = 0.1% formic acid in acetonitrile). The top 15 most intense precursor ions from each full scan (resolution 120,000) were isolated for HCD MS2 (resolution 15,000, HCD Collision Energy 30%) with a dynamic exclusion time of 60 s, a target AGC of 1E5 and a maximum injection time of 50 ms. Precursors with 3+ to 6+ charge states were included. Each sample was analyzed a second time using the same parameters, except that 2+ precursors were also included for HCD MS2 to find more SUMOylated peptides. We prepared two samples from the wild type and the *slx8-1* mutant, respectively. This dataset consisted of 24 raw files. The workflow for the dataset preparation is illustrated in **Fig. 2A**.

**Fig. 2.**
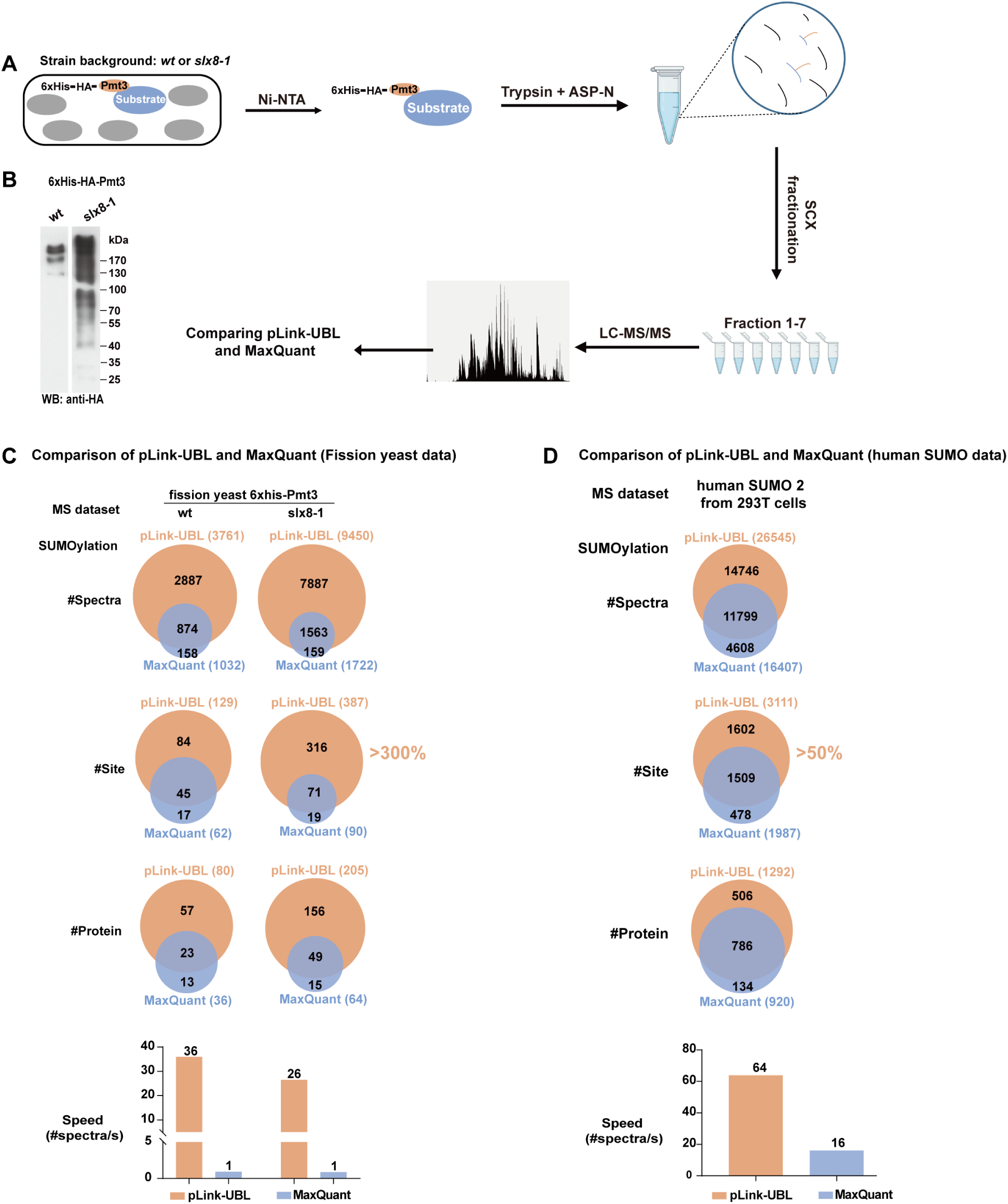
Identification of SUMOylation sites on fission yeast proteins using pLink-UBL, in comparison to MaxQuant. **(A)** Sample preparation and data collection from profiling of protein SUMOylation sites in fission yeast. 6×His-HA-Pmt3 was expressed in the WT background as well as the Slx8-1 temperature-sensitive mutant background to accumulate more SUMOylated proteins. **(B)** Western blot analysis of nickel-beads enriched 6×His-HA-Pmt3 protein from WT or Slx8-1 mutant cells. Comparison of pLink-UBL and MaxQuant on identification of Pmt3/SUMO modified peptides. **(C)** Comparison of pLink-UBL and MaxQuant on re-identification of peptides modified by human SUMO2, using published dataset [28].

Database search was performed with Asp-N and trypsin specificity and five missed cleavage sites were allowed. Peptides were accepted with a length range between 6 and 60 amino acids and a mass range between 600 and 6,000 Da. Deamidated[N], Carbamidomethyl[C], Carbamyl[AnyN-term], and Oxidation[M] were chosen as variable modifications. Mass tolerances for the precursor and the fragment were both set to 20 ppm to search against the yeast protein database (5,251 proteins). For speed evaluation, search engines used eight threads for the search, and the computing times were measured on the same computer equipped with (Windows Server, Intel Xeon E5-2670 CPU with 32 cores, 2.6 GHz, 128 GB RAM).

### Preparation of the Human SUMO2 dataset and database search

The Human SUMO2 dataset is from a published human cell proteome dataset [28], which consisted of 12 raw files.

Database search was conducted using Lys-C and Asp-N specificity, with the allowance of five missed cleavage sites. Peptides within the length range of 6 to 60 amino acids and a mass range of 600 to 6000 Da were considered. Carbamidomethyl[C] was chosen as a fixed modification, and Acetyl[ProteinN-term] and Oxidation[M] were chosen as variable modifications. Mass tolerances for precursor and fragment were both set to 20 ppm to search against the Human protein database (92,952 proteins). For speed evaluation, two search engines utilized fifteen threads for the search (Windows, AMD Ryzen 7 3800X with 16 cores, 3.9 GHz, 32 GB RAM).

## Results

### Development of pLink-UBL for identification of UBL modification sites

For identification of UBL modification sites using the conventional PTM search strategy **(Fig. 1A and B, strategy 1)**, it is necessary to mutate a residue close to the C-terminus to create a trypsin cleavage site [32, 33]. However, sequence information unique to each UBL is lost or greatly reduced in the resulting tryptic peptide. Further, there is a possibility that mutation may compromise the function of a UBL.

Alternatively, a relatively long C-terminal peptide of UBL could be left attached to a substrate peptide and these two peptides could be identified as a pair of cross-linked peptides linked through an isopeptide bond. Such cross-linked peptide pairs can be identified from their MS2 spectra by cross-link search engines such as pLink [37, 38]. However, for best results, such search engines need to be optimized for UBL cross-links. Therefore, based on the framework of pLink, we started to develop pLink-UBL.

For software training, we generated two UBL cross-linked peptide libraries. A total of 246 substrate peptides were synthesized in groups of 6 or 9 peptides **(Supplementary Fig. S4)**, which are all tryptic peptides containing an internal lysine residue. Two UBL C-terminal peptides were synthesized: DQIEAVLEQLGG (Pmt3-C-pep) and DVFQQQTGG (SUMO2-C-pep), which represent, respectively, the C-terminal remnant peptide of fission yeast SUMO Pmt3 and that of human SUMO2 after trypsin and Asp-N digestion. Each UBL peptide was chemically linked to each of the 246 substrate peptides. To facilitate formation of the isopeptide bond, all the reactive groups were protected except the C-terminal carboxylate of UBL-C-pep and the [-amine of the internal lysine in substrate peptides. The protective groups were removed after the cross-linking reaction.

Liquid chromatography-tandem mass spectrometry (LC-MS/MS) analysis of the two synthetic UBL-substrate peptide libraries produced two training datasets, Syn-Pmt3 and Syn-SUMO2, consisting of 1970 and 1560 high-quality, non-redundant MS2 spectra, respectively. They contain fully annotated HCD (higher-energy collisional dissociation) spectra selected from the raw MS data **(Fig. 1C)**. Analysis of these datasets revealed fragmentation characteristics of these peptides **(Supplementary Fig. S2)** and the significance rank of different ions types **(Supplementary Fig. S5 and S6),** definition of ions types can be found in **Supplementary Fig. S7**. After multiple rounds of evaluation, a total of 18 ion types are included in the fine-scoring algorithm of pLink-UBL **(Supplementary Fig. S8, note1)**

**Fig. 1D** illustrates the workflow of pLink-UBL. First, MS data are processed in the same way as pLink, *i.e.,* using pParse [54] to detect the monoisotopic peaks in MS1 and to subsequently calibrate the precursor mass for each MS2. Then, UBL-linked substrate peptides are identified in two stages. Since the search space of UBL peptides (a limited number of UBL proteins) is much smaller than that of substrate peptides (entire proteome for a given species), pLink-UBL

identifies UBL-C-pep before it identifies the cross-link between a UBL-C-pep and a substrate peptide **(see Supplementary note 2 for details).** As shown in **Supplementary Fig. S9**, the estimated FDR by pLink-UBL is close to or slightly higher than the true FDR values over the (∼5%) range.

### Performance evaluation of pLink-UBL

We tested pLink-UBL in three different ways by using datasets of simulated MS2 spectra, synthetic UBL-linked peptides, and those of real-world samples.

Two simulated test datasets Sim-Pmt3-Ecoli and Sim-SUMO2-Ecoli were generated using pSimXL [37]. Each consists of 5000 simulated HCD spectra generated from the sequences of 1000 *E. coli* proteins for substrate peptide, plus the C-terminal peptide sequence of either Pmt3 or Human SUMO 2. Each simulated dataset also contains 5000 HCD spectra of linear peptides. These spectra, 10,000 in total for each dataset, were paired with correct precursor mass values as well as incorrect ones by introducing a +5 or + 10 Da mass shift to serve as negative data.

The two datasets of synthetic UBL-linked peptides consist of the positive data (Fig. 1C) and equal numbers of HCD spectra of unmodified linear peptides as the negative data. The search space was intentionally increased or not, by adding the human protein database to mimic real-world situations.

Using the above datasets, we compared pLink-UBL to pLink and pFind 3. As shown in **Fig. 1E-H**, pLink-UBL and pLink displayed 100% sensitivity and 95.8%-100% precision across all the datasets tested. The sensitivity of pFind 3—representing the PTM search strategy dropped severely as the search space increased to include the human protein database, to as low as 77.5%.

The original pLink, although performed well in terms of sensitivity and precision, incurred a time cost 3-23 times higher than that of pLink-UBL.

Next, we evaluated pLink-UBL against MaxQuant using real-world data. To generate test data, we expressed 6×His-tagged Pmt3 in fission yeast, in a wild-type strain as well as a *slx8-1* strain **(Fig. 2A)**. Slx8 is a SUMO-dependent ubiquitin ligase, deletion of which allows SUMOylated proteins to accumulate to a higher level [55]. This is verified by western blotting **(Fig. 2B)**. 6×His-Pmt3 protein samples enriched with nickel beads were digested using trypsin and Asp-N and fractioned before LC-MS/MS analysis. At 5% FDR for cross-link-spectrum match (CSM) and a minimum of two spectra for each identification, the number of Pmt3-modified peptides identified from *wild*-type and *slx8-1* cells are 251 and 495 by pLink-UBL and 94 and 120 by MaxQuant, respectively. The number of corresponding spectra and Pmt3-modified sites are shown in **Fig. 2C** (see **Supplementary Tables S3 and S4** for details). Taking the Slx8-1 data as an example, MaxQuant identified 1722 spectra and 90 sites, pLink-UBL identified 9450 spectra and 387 sites, with an overlap of 1563 spectra and 71 sites. In other words, 90.7% of the spectra and 78.8% of the sites identified by MaxQuant, were identified by pLink-UBL. pLink-UBL additionally had many unique identifications, about five times as many as the overlap.

From the *wild*-type and *slx8* mutant strains, we identified a total of 459 fission yeast SUMOylation sites. A previous study identified 1028 sites in fission yeast using a mutagenesis strategy shown in **Fig. 1B (upper right).** Although fewer SUMOylation sites are identified in this study, it is certain that there are no misidentified ubiquitination sites in them, because the 12-aa remnant peptide of Pmt3 is easily distinguishable from the diglycyl remnant of ubiquitin. Motif analysis showed that the same motifs KXE and (D/E)XK, where K is the SUMOylated lysine, and X is any amino acid, are enriched among the sequences identified in both studies. **(Supplementary Fig. S10).**

Lastly, we compared pLink-UBL and MaxQuant on a published dataset of human

SUMO2/3-modified peptides [28]. This dataset originated from a Lys-C digested HEK 293T cell lysate, in which the C-terminal peptide of endogenous, wild-type SUMO2/3 and its covalently linked substrate peptides were enriched using an immobilized antibody. As shown in **Fig. 2D**, pLink-UBL increased the number of identified SUMOylation sites by 57%, from 1987 to 3111 (see **Supplementary Tables S5 and S6** for details), and the search speed of pLink-UBL was 4 times faster than MaxQuant. Taken together, we demonstrate that pLink-UBL has a clear advantage in sensitivity, precision, and speed in UBL modification site analysis.

### Identification of protein Neddylation and Urmylation sites in fission yeast using pLink-UBL

Having validated pLink-UBL on analysis of SUMOylation sites, we used it to map modification sites of two additional fission yeast UBLs Ned8 and Urm1. Samples were prepared and analyzed in the same way as for Pmt3 **(Fig. 3A).** WB analysis of the Ni-NTA enriched samples showed that, in contrast to Pmt3, which displayed many Pmt3-positive bands from low to high molecular weight, Ned8 and Urm1 positive bands appeared to be rather limited **(Fig. 3B)**. Consistently, pLink-UBL search of the corresponding LC-MS/MS data identified a handful of modification sites for Ned8 and Urm1, compared to hundreds for Pmt3/SUMO **(Fig. 3C)**. Specifically, 5 Neddylation sites were identified, 4 on Ned8 itself and 1 on Cul4. All five Neddylation sites have been previously published [34, 56, 57] and are supported by high-quality MS2 spectra **(Supplementary Fig. S11)**. This finding demonstrates that pLink-UBL can be utilized to identify additional UBL modifications.

**Fig. 3.**
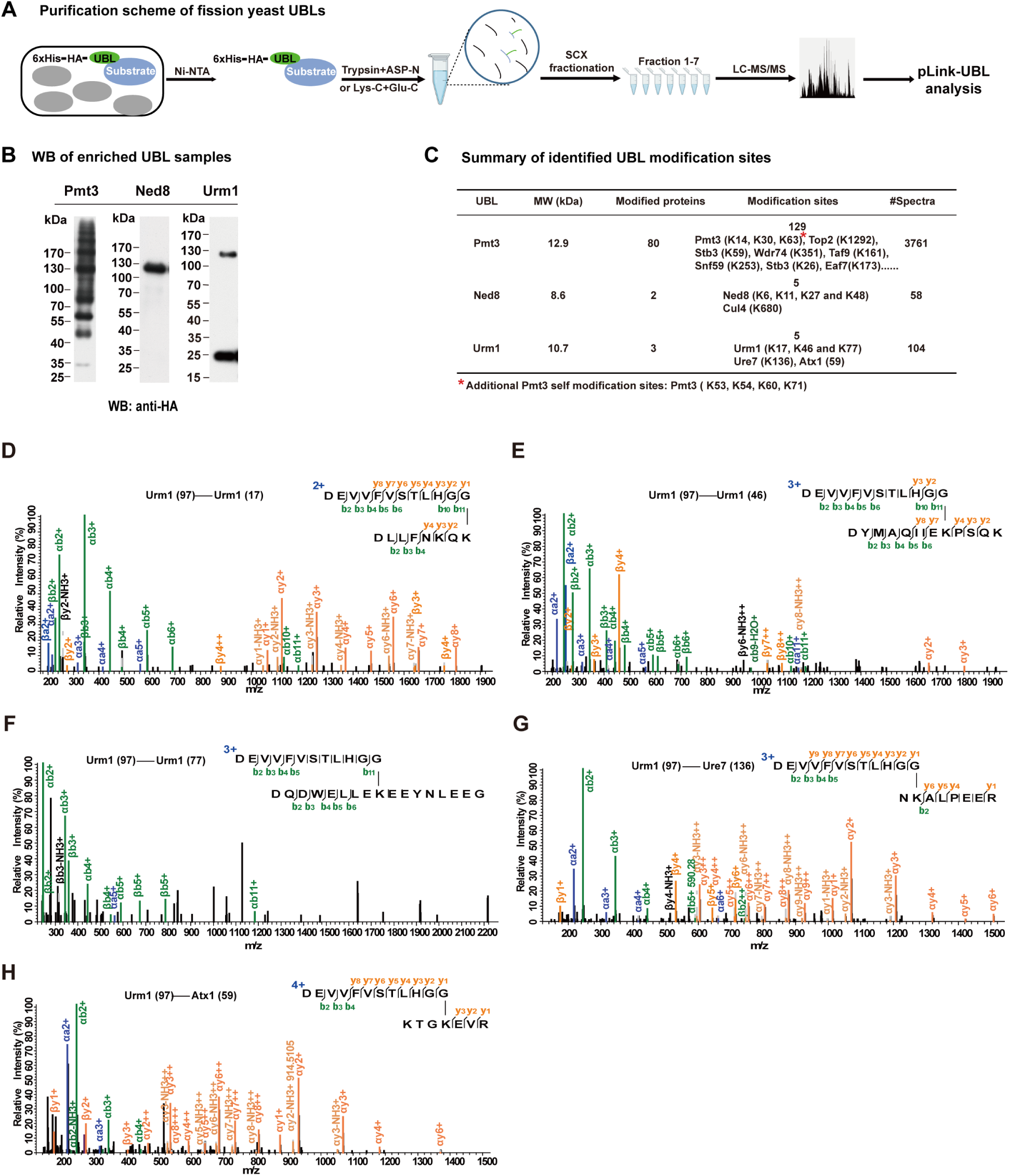
Identification of protein Neddylation and Urmylation sites using pLink-UBL. **(A)** Procedure for identification of UBL modification of fission yeast proteins**. (B)** Western blot analysis of nickel-beads enriched Ned8 and Urm1 proteins, with Pmt3 served as a reference. **(C)** Summary of the identification results, with the Pmt3 result listed as a reference. **(D-H)** MS2 spectra of five Urm1-modified peptides/sites identified by pLink-UBL.

Likewise, 5 Urmylation sites were identified, 3 on Urm1 itself, 1 on Ure7, and 1 on Atx1. These Urm modification sites were not known previously. We examined their MS2 spectra manually. Except for Urm1(77), which has a median-quality MS2, the other four sites-Urm1(17), Urm1(46), Ure7(136), and Atx1(59) all have high-quality MS2 **(Fig. 3D-H)**.

Like Ub, Pmt3/SUMO, Ned8, and Urm1 all modify themselves, with 7, 4, and 3 self-modification sites identified from wild-type cells, respectively **(Fig. 3C)**. Two additional Pmt3 self-modification sites (K39, K51) were found in slx8 mutant cells. Previous studies have made similar observations [28, 32–34]. This suggests that, it may be common for UBLs to form chain structures, just like ubiquitin.

### Identification of small molecule substrate of Pmt3/SUMO

Since the C-termini of Ub and certain UBLs such as ATG8 and LC3 can be conjugated to small molecules, we wondered whether this could happen to other UBLs. We focused on Pmt3/SUMO as it is of high abundance in fission yeast. To enrich possible small molecule modifications at the C-terminus of Pmt3, we developed rabbit polyclonal antibodies against a 27-aa peptide of Pmt3 (IG27) whose sequence corresponds to the most C-terminal peptide of Pmt3 after trypsin digestion **(Supplementary Fig. S12A and Methods).** The anti-IG27 antibody (#2517) exhibited high specificity and high efficiency towards enrichment of IG27, either as a free peptide, in a pair of cross-linked peptides, or as a segment of a fusion protein **(Supplementary Fig. S12B-D).**

Using the anti-IG27 antibody, we developed a workflow to look for non-protein substrates of Pmt3/SUMO. As shown **(Fig. 4A)**, this workflow starts with Ni-NTA enrichment of 6×His-tagged Pmt3 from fission yeast homogenates. This is followed by trypsin digestion and enrichment of the C-terminal peptide of Pmt3, either modified or unmodified, using the anti-IG27 antibody immobilized on protein A agarose beads. After LC-MS/MS analysis of the enriched peptides using a high-resolution Q-Orbitrap instrument, pFind blind search is used to look for peptides with unexpected modifications.

**Fig. 4.**
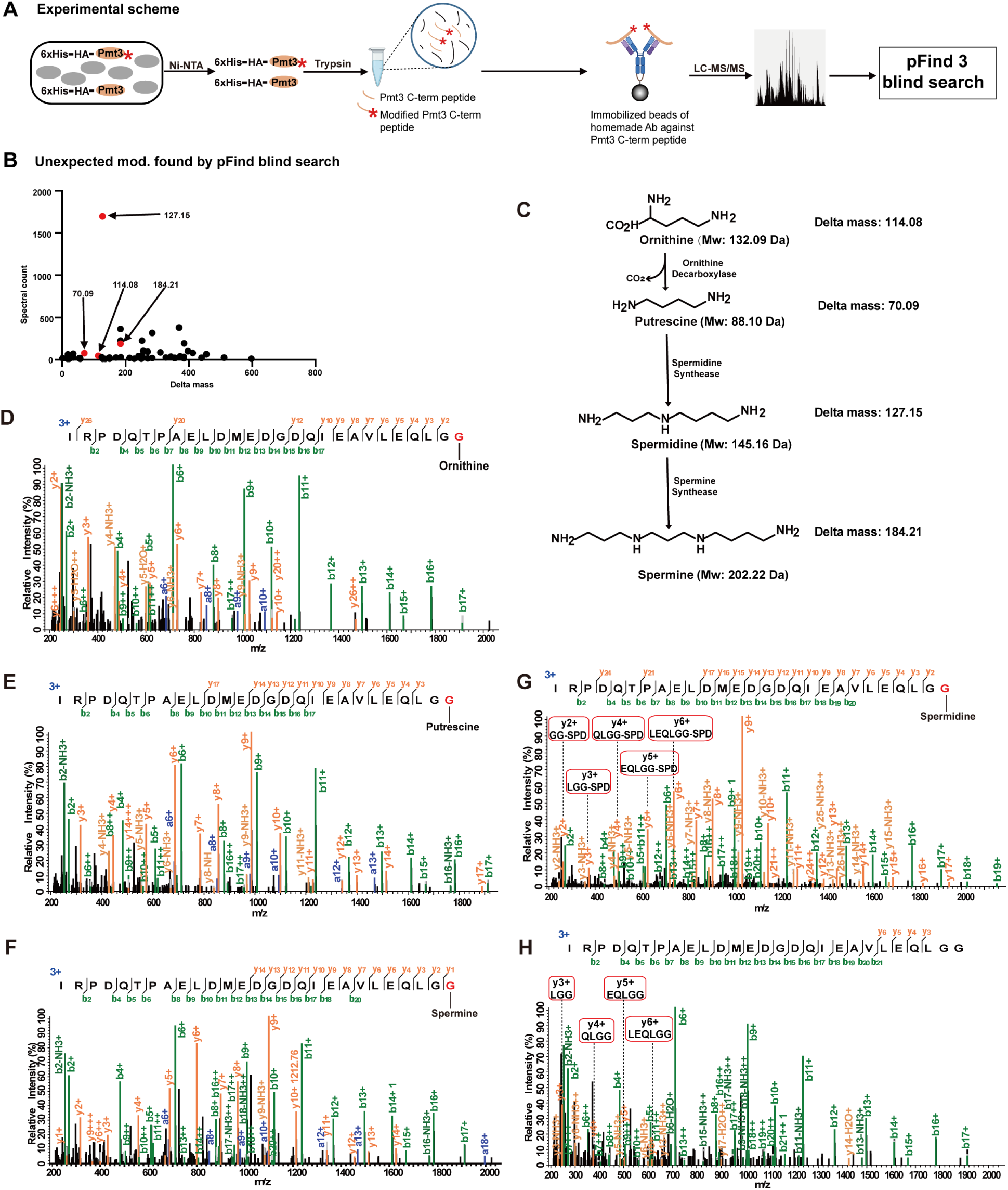
Identification of small molecule substrates of Pmt3/SUMO. **(A)** Experimental scheme. A rabbit antibody was generated, which can bind to the 27-aa C-terminal peptide of Pmt3 with high affinity. This antibody was used to enrich Pmt3 C-terminal peptides following the enrichment of 6×His-HA-Pmt3 protein. The resulting MS data were searched using pFind 3 under the blind search mode to identify unexpected modifications. **(B)** Frequency of unexpected modifications found by pFind 3 blind search. A total of 57 distinct delta mass values were reported from the Pmt3 sample generated in (A). **(C)** The polyamine biosynthesis pathway. **(D-G)** MS2 of the Pmt3 C-terminal peptide with putative modification of ornithine, putrescine, spermine, and spermidine, respectively. **(H)** MS2 of unmodified Pmt3 C-terminal peptide.

The frequencies of unexpected modifications detected in the sample of Pmt3 C-terminal peptide, expressed as Δmass values, are displayed in **Fig. 4B**. Out of the 57 unexpected modifications on the C-terminal peptide of Pmt3, Δmass of 127.15 stands out with 1698 supporting MS2. Looking into the Human Metabolome Database (HMDB), we found only one entry that matches with Δmass of 127.15 with a mass tolerance of 0.0005 Da. This is spermidine **(Fig. 4C).**

Spermidine is a polyamine with important biological functions in protein synthesis, aging, autophagy, and DNA stabilization [58–61]. It is generated from putrescine, which in turn is generated from ornithine **(Fig. 4C)**. And spermidine can be converted further into spermine **(Fig. 4C)**. Interestingly, Δmass values that correspond to the three other polyamine modifications were also reported by pFind blind search **(Fig. 4B).** MS2 spectra are able to localize the putative modification of Pmt3/SUMO by ornithine, putrescine, spermine, and spermidine to the last 1-3 amino acids **(Fig. 4D-G)**. This is demonstrated by a head-to-head comparison of the MS2 spectra of the Pmt3 C-terminal peptide with or without spermidine modification **(Fig. 4G and H)**. Since the last three amino acids of Pmt3 are LGG-COOH, which contain no chemically reactive side chains, it leaves the very C-terminal carboxylate group as the only reasonable site for polyamine modification.

### The C-terminus of Pmt3/SUMO is conjugated to the N^1^ or also likely, N^8^ position of spermidine

Spermidine has two primary amino groups, N^1^ and N^8^ **(Fig. 4C)**. To find out whether only one of them, or both, can be attached to the C-terminus of Pmt3, we examined the fragmentation spectrum of free spermidine **(Fig. 5A)** and those of Pmt3-C-pep-Spd **(Fig. 5B-F)**.

**Fig. 5.**
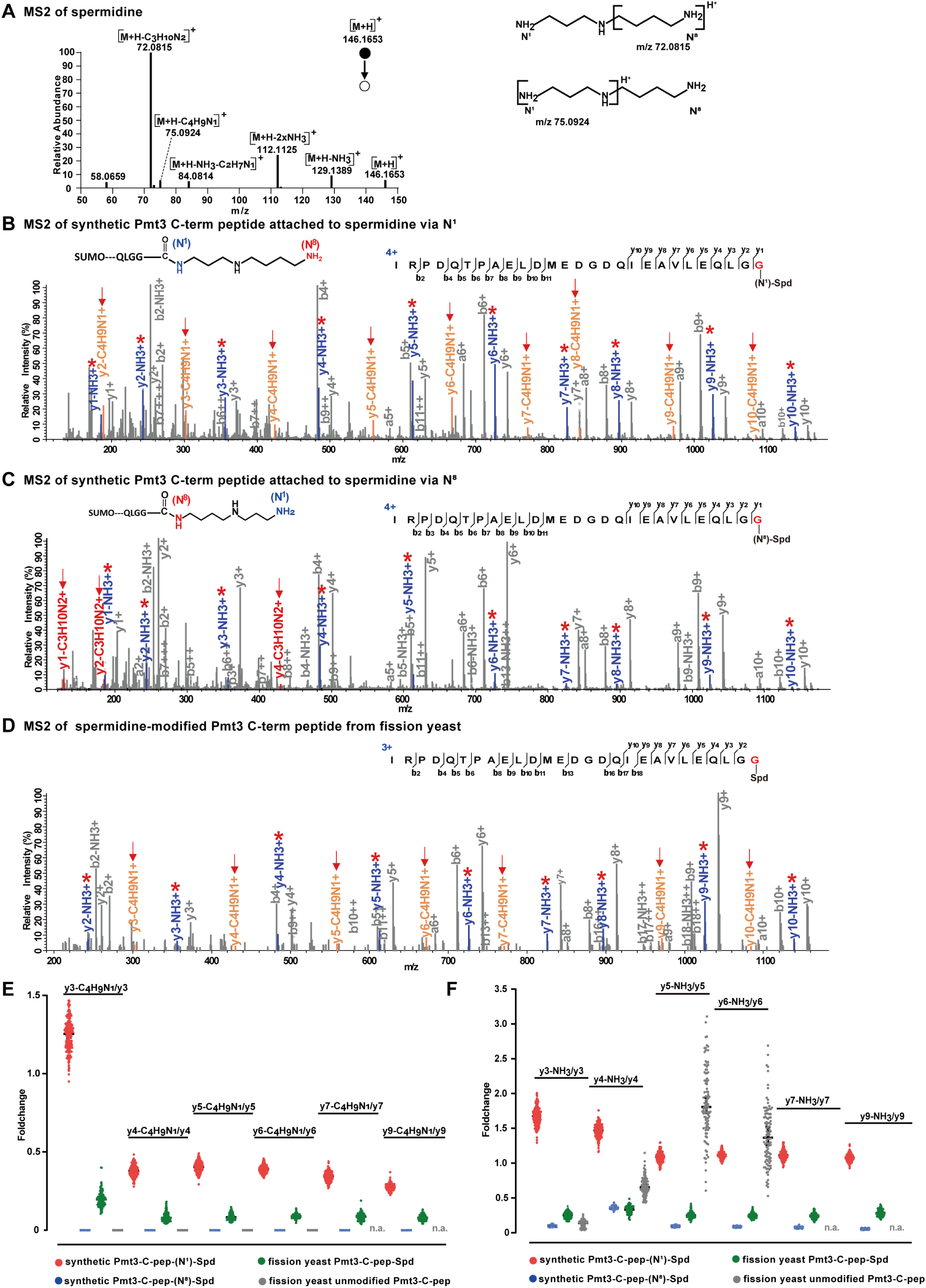
The C-terminus of Pmt3/SUMO is conjugated to spermidine via the N^1^ position, and possibly also at the N^8^ position. **(A)** Annotated MS2 of free spermidine. Cleavages that generate m/z 72.0815 and m/z 75.0924 are proposed and shown on the right. **(B)** MS2 of chemically synthesized Pmt3-C-pep-(N^1^)-spermidine. Note the characteristic 71.0375 Da (-C4H9N1) NL ions and the high abundance of NH3 NL ions. **(C)** MS2 of chemically synthesized Pmt3-C-pep-(N^8^)- spermidine. Note the absence of characteristic fragment ions and the low abundance of NH3 NL ions. **(D)** MS2 of Pmt3-C-pep-spermidine from fission yeast. Note the presence of 71.0375 Da NL ions and NH3 NL ions, but their intensities are lower than their counterparts in (B). **(E and F)**. Intensity of y ions with 71.0375 Da NL or NH3 NL nominalized to their cognate y ions. The statistics are based on 200 MS2 of synthetic Pmt3-C-pep-(N^1^)-spermidine, 200 MS2 of synthetic Pmt3-C-pep-(N^8^)-spermidine, 128 MS2 of Pmt3-C-pep-spermidine from fission yeast, and 100 MS2 of unmodified Pmt3-C-pep from fission yeast. 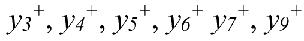 are counted for all except the unmodified Pmt3-C-pep, which lacks 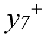 and 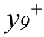 ions. Note that as for the normalized intensity values, Pmt3-C-pep-spermidine comes between Pmt3-C-pep-(N^1^)-spermidine and Pmt3- C-pep-(N^8^)-spermidine. For unmodified Pmt3-C-pep, the relative intensity of 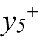-NH3 and 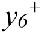- NH3 are erratically high, probably because denominator 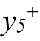 and 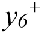 ions are of very low intensity.

We collected MS2 spectra of IG27-(N^1^)-Spd and IG27-(N^8^)-Spd, which are the synthetic Pmt3 C-terminal peptide with covalently linked spermidine through N^1^ and N^8^, respectively. Side-by-side comparison of the MS2 spectra shows that IG27-(N^1^)-Spd produces a characteristic neutral loss of 71.0735 Da off the *y^+^* ions detected, from *y_2_^+^* to *y_10_^+^*, and the high abundance of NH_3_ NL ions **(Fig. 5B, C and F)**. The MS2 of spermidine-modified Pmt3 C-terminal peptide from fission yeast samples matched with that of IG27-(N^1^)-Spd, with the characteristic NL 71.0735 peaks associated with the *y* ions. However, the intensity of *y* ions with 71.0375 Da NL or NH_3_ NL are lower than their counterparts in (B) **(Fig. 5B-F)**. Based on these results, we propose that in fission yeast the C terminus of Pmt3/SUMO is conjugated to the N^1^ or also likely, N^8^ position of spermidine.

### Reversible Pmt3/SUMO-spermidine conjugation is catalyzed by enzymes

Next, we asked whether SUMO-spermidine conjugation forms spontaneously or is catalyzed enzymatically. In fission yeast, the SUMO E1, E2, and E3 enzymes are Rad31/Fab2, Hus5, and Pli1, respectively **(Fig. 6A)**. Together, they are responsible for conjugating Pmt3/SUMO to protein substrates. We wondered whether they are required for conjugating Pmt3/SUMO to spermidine. We expressed and purified recombinant fission yeast SUMO, E1, E2, and E3 to set up an *in vitro* assay. As shown in **Fig. 6B**, E1 (Rad31/Fab2), E2 (Hus5), and ATP are required for the formation of a product band below Pmt3/SUMO. In-gel digestion and LC-MS/MS analysis of this product band revealed that it is Pmt3 with spermidine attached to its C-terminus **(Fig. 6C)**. This assay also showed that whereas E1, E2, and ATP are required for Pmt3-spermidine ligation, E3 (Pli1) is not.

**Fig. 6.**
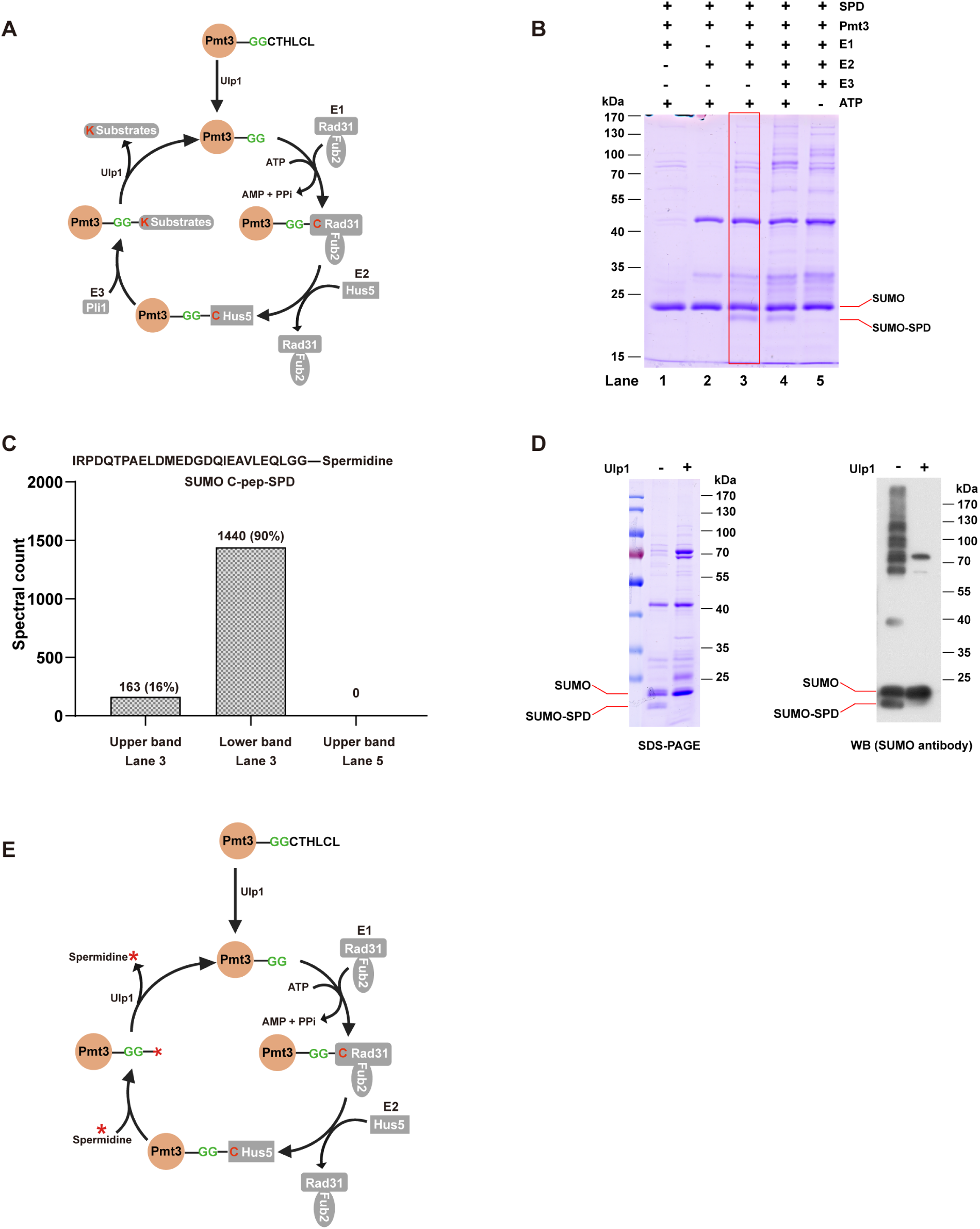
Pmt3/SUMO-spermidine conjugation requires ATP, E1, and E2, but not E3, and can be reversed by SUMO isopeptidase Ulp1. **(A)** Conjugation mechanism of SUMO to protein substrates in fission yeast. **(B)** In vitro assay of Pmt3-spermidine conjugation. **(C)** LC-MS/MS analysis result of Pmt3/SUMO from (B). **(D)** In vitro separation of Pmt3 from covalently linked spermidine by Ulp1. **(E)** Proposed mechanism for SUMO-spermidine conjugation and separation.

In fission yeast, the SUMO protease Ulp1 is responsible for processing newly synthesized Pmt3/SUMO into its mature form by cleaving off the last 6-aa tail, to expose the diglycine C-terminus **(Fig. 6A)**. Ulp1 is also responsible for deSUMOylation by cleaving the isopeptide bond formed between the SUMO C-terminal carboxylate and a lysine side-chain amine of a substrate protein. Our in vivo assay showed that Ulp1 can also remove Pmt3/SUMO from the spermidine attachment **(Fig. 6D)**. Based on the above results, we propose a mechanism for reversible Pmt3-spermidine conjugation that involves SUMO E1, E2, and Ulp1 **(Fig. 6E)**. SUMO E3 is not part of this mechanism.

### SUMO-Spermidine conjugation in budding yeast, mouse, and human cells

Wondering whether SUMO-spermidine conjugation is a general phenomenon, we collected published SUMO proteomics data of mouse and human cells. We selected the ones that enabled detection of the SUMO C-terminal peptide of sufficient length (>8 aa), this is, from trypsin or trypsin+Asp-N digested wild-type SUMO protein samples. A variable modification search using pFind 3 found MS2 spectra of SUMO-spermidine in the proteomics data of mice and humans **(Fig. 7A).** The spectral count percentage of SUMO-spermidine over the summed total of SUMO and SUMO-spermidine, concerning only the C-terminal peptide, reached 18% for mouse and 2% for human cells. Representative MS2 spectra of SUMO-spermidine for the two species are shown in **Fig. 7B-C**. Similar results were obtained when additional datasets were analyzed **(Supplementary Fig. S13)**. Based on the above evidence, we propose that SUMO-spermidine conjugation occurs widely in eukaryotes, from yeast cells to humans.

**Fig. 7.**
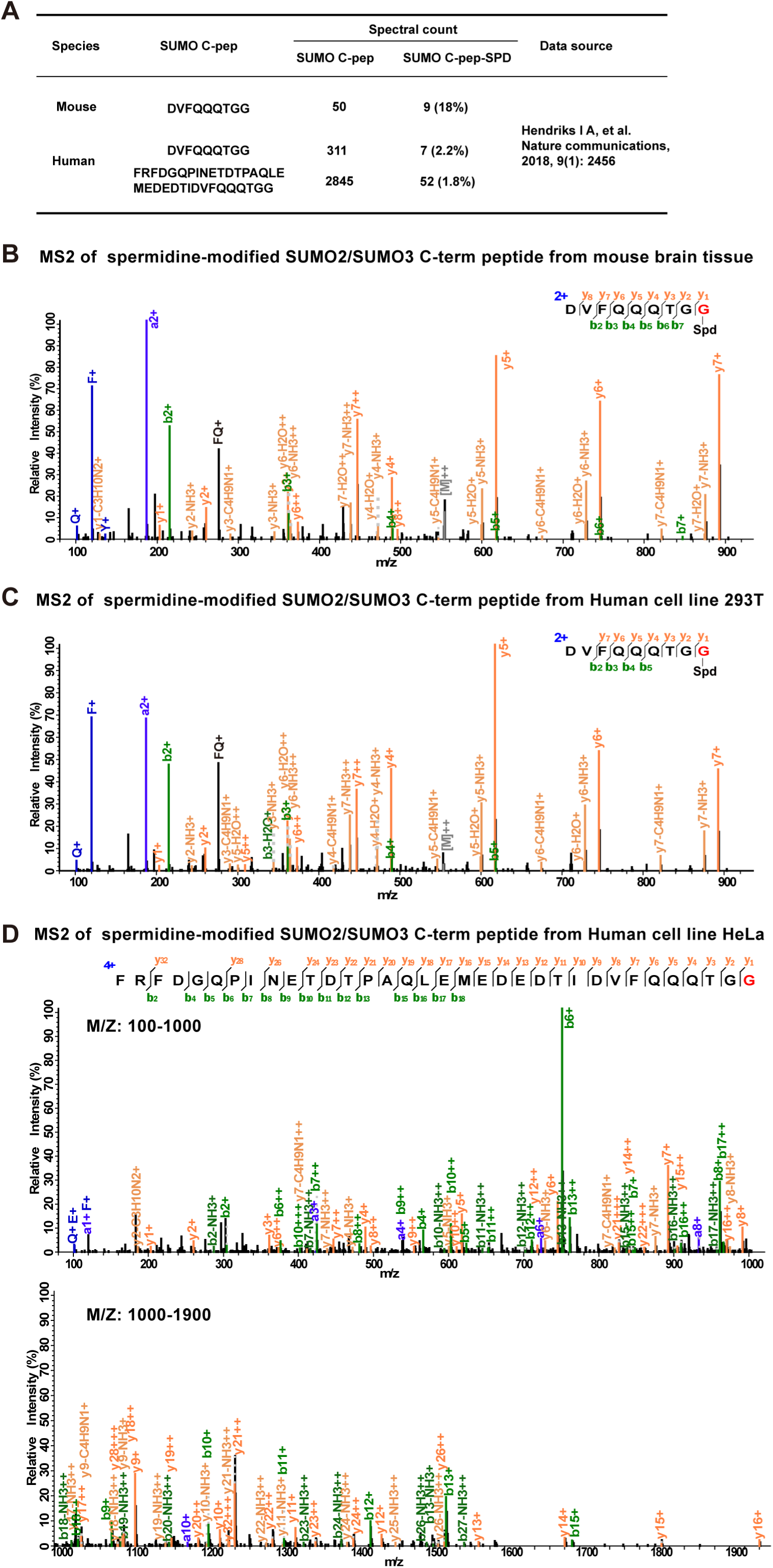
Evidence of SUMO-spermidine conjugation in mice and humans. **(A)** SUMO- spermidine conjugation identified by reanalysis of published proteomics data of mouse and human origins. **(B, C, and D)** Representative MS2 of mouse (B) and human (C and D) SUMO2/3 C-terminal peptide with spermidine attached at the C terminus.

### Ubiquitin can be conjugated to spermidine in the presence of E1, E2, and ATP

Having discovered that SUMO-spermidine conjugation is a common biochemical reaction in eukaryotic cells catalyzed by E1 and E2 with energy supply from ATP, we expanded our investigation to ubiquitin of the fission yeast. Unlike SUMO, which has a single E1 and a single E2, Ub has one E1 and 11 E2 enzymes in fission yeast. We started the *in vitro* assay with Uba1 and Ubc4, which is one of the 11 E2 enzymes. As shown in **Fig. 8A**, the Ub band was up-shifted when E1, E2, and ATP were all added to the reaction. Lys-C digestion which generates a 13-aa C-terminal peptide of Ub, followed by LC-MS/MS analysis, found that out of the 1820 MS2 spectra that matched with the C-terminal peptide of Ub, 1570 (86.3%) were of Ub-spermidine conjugation **(Fig. 8B).**

**Fig. 8.**
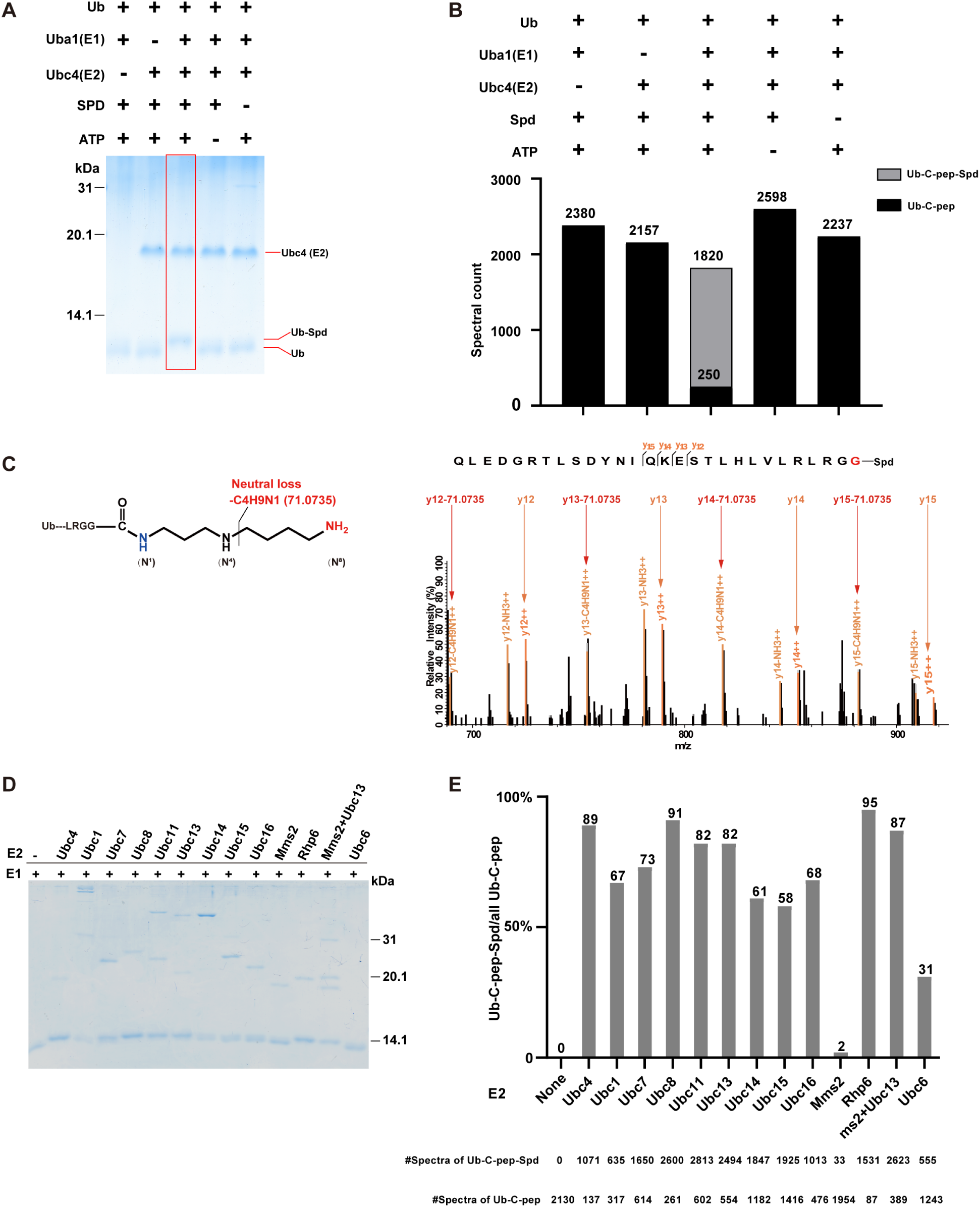
In vitro conjugation of Ub and spermidine in the presence of E1, E2, and ATP. **(A)** In vitro assay of Ub-spermidine conjugation. **(B)** LC-MS/MS analysis results of Ub from reaction under different conditions in (A). **(C)** Annotated MS2 of Ub-spermidine. For clarity, only the m/z range (680-920) is shown. **(D)** In vitro conjugation of Ub-spermidine by E1 and different E2 enzymes. **(E)** LC-MS/MS analysis results of the Ub protein band from the different reactions in (D).

MS2 spectra indicate that Ub-spermidine conjugation occurs also at the N^1^ position of spermidine, judging by the high abundance of neutral loss of 71.0735 Da **(Fig. 8C)**.

Ub-spermidine conjugation at the N^8^ position of spermidine is a possibility that we can neither confirm nor reject.

Lastly, we expressed and purified successfully 12 fission yeast E2 proteins of Ub and tested whether they could conjugate Ub to spermidine. Of the 12 E2 proteins, Mms2 and Ubc13 form a heterodimeric E2 enzyme in which Ubc13 is the catalytic subunit, and Mms2 is a regulatory subunit. On SDS-PAGE, all of the tested E2 proteins up-shifted the protein band of Ub except Mms2, which failed to up-shift Ub to the same degree as the others **(Fig. 8D)**. LC-MS/MS results indicated that all except Mms2 can conjugate this reaction *in vitro*.

In conclusion, we find that spermidine can be conjugated to Ub and UBL, and Ub/UBL-polyamine conjugation may be common across eukaryotic species.

## Discussion

This study aims to profile SUMOylation sites in a precise and comprehensive manner. To reach this goal, we designed a strategy in which no mutation was introduced to the SUMO protein, developed pLink-UBL to improve data analysis, and generated a high-affinity antibody to further enrich the C-terminal peptide of Pmt3/SUMO after enrichment of the Pmt3/SUMO protein. However, only 67 Pmt3 modification sites were identified using this strategy. This is far less than what had been identified using the antibody-free procedure shown in **Fig. 2A** and **Fig. 3A**. Sample loss during the 2-step enrichment procedure **(Fig. 4A)** is probably to blame; the amount of twice enriched sample was so limited that further fractionation did not improve identification. Nevertheless, the number of Pmt3-modified proteins identified in this study (200+) is on par with an earlier report [62].

Over a thousand Pmt3 modification sites were reported in a 2015 study [32], which relied on the mutagenesis strategy shown in **Fig. 1B (strategy 1)**. The number of Pmt3 modification sites identified in this study did not exceed 500 in total. The large gap between the two numbers may have to do with the differences in the method and the number of experimental conditions analyzed. This study analyzed only *wt* and the *slx8* mutant; the 2015 study examined more mutants. The method used in this study can distinguish SUMOylation from ubiquitination. In contrast, the mutagenesis-based method used in previous studies cannot. This would lead to false positive identifications if SUMOylated proteins are also ubiquitinated. As such, the actual number of SUMOylation sites is probably lower than what is detected.

Spermidine, a prominent member of polyamines, can bind to RNA and DNA [63–65]. When spermidine is attached to a lysine residue (K50 or K51) of eukaryotic translation initiation factor 5A (eIF5A), it can lead to hypusination of eIF5A under the actions of specialized enzymes. Hypusination activates eIF5A and is required for cells to grow [59, 61, 66]. Spermidine can also be attached to glutathione, a tripeptide antioxidant [67]. However, Ub/UBL-spermidine conjugation was not previously known. A chemical biology approach to search for spermidine-binding proteins using a functionalized spermidine analog found 140 proteins, but Ub and UBL are not among them [68]. Here we show that SUMO-spermidine conjugation is common among eukaryotes from fission yeast to humans, and formation of SUMO-spermidine requires E1, E2, and ATP, but not E3. This finding originated from antibody enrichment of the C-terminal peptide of SUMO, followed by LC-MS/MS analysis and pFind 3 blind search. Such a strategy should be effective elsewhere, if the goal is to look for unexpected small molecule

modifications of proteins.

## Supplementary Data

This article contains Supplementary data [37, 38, 69]

## Author Contributions

Meng-Qiu Dong and Li-Lin Du devised this project. Guang-Can Shao and Shan-Lu carried out the MS experiments, yeast experiments, the development of peptide antibody, and data analysis. Zhen-Lin Chen and Si-Min He developed and evaluated the software pLink-UBL, Xiang-Bing Qi and Qing-Cui Wu synthesized SUMOylated peptides. Hao Chi instructed with pFind blind search. Yao Sheng, Jing Wang, and Jian-Hua Sui carried out the experiments on the immunization of the rabbit. Guang-Can Shao, Zhen-Lin Chen, Li-Lin Du, and Meng-Qiu Dong wrote the manuscript.

## Conflict of interest

All the authors declare that they have no conflict of interest with the contents of this article.

## Acknowledgments

The authors would like to thank Dr. Yong Cao for helpful discussions. This work was funded by intramural grants from NIBS, Beijing and TIMBR, Tsinghua University.

## Data availability

The mass spectrometry proteomics data have been deposited to the ProteomeXchange Consortium (https://proteomecentral.proteomexchange.org) via the iProX partner repository [70, 71] with the dataset identifier PXD058355 (Username: shaoguangcan; Password: donglab_NIBS).

## Abbreviations

DDA: Data-Dependent Acquisition
HCD: Higher-Energy Collision-Induced Dissociation
IAA: Iodoacetamide
LC-MS/MS: Liquid chromatography-tandem mass spectrometry
OD: Optical Density
PTM: Post-Translational Modification
SCX: Strong Cation Exchange
TCA: Trichloroacetic Acid
UBL: Ubiquitin Like Protein
YES: Yeast Extract with Supplement

## Notes

### Competing Interest Statement

The authors have declared no competing interest.

## References

1. 1. van der Veen, A.G. and H.L. Ploegh. (2012) Ubiquitin-like proteins. Annu Rev Biochem. 81, 323–357.

2. Hochstrasser, M. (2009) Origin and function of ubiquitin-like proteins. Nature. 458(7237), 422–429.

3. Hochstrasser, M. (2000) Evolution and function of ubiquitin-like protein-conjugation systems. Nat Cell Biol. 2(8), E153–157.

4. Vijay-Kumar, S., C.E. Bugg, and W.J. Cook. (1987) Structure of ubiquitin refined at 1.8 Åresolution. J Mol Biol. 194(3), 531–544.

5. Schulman, B.A. and J.W. Harper. (2009) Ubiquitin-like protein activation by E1 enzymes: the apex for downstream signalling pathways. Nat Rev Mol Cell Biol. 10(5), 319–331.

6. Michelle, C., P. Vourc’h, L. Mignon, and C.R. Andres. (2009) What was the set of ubiquitin and ubiquitin-like conjugating enzymes in the eukaryote common ancestor? J Mol Evol. 68, 616–628.

7. Jentsch, S., W. Seufert, T. Sommer, and H.A. Reins. (1990) Ubiquitin-conjugating enzymes: novel regulators of eukaryotic cells. Trends Biochem Sci. 15(5), 195–198.

8. Buetow, L. and D.T. Huang. (2016) Structural insights into the catalysis and regulation of E3 ubiquitin ligases. Nat Rev Mol Cell Biol. 17(10), 626–642.

9. Walsh, C.K. and A. Sadanandom. (2014) Ubiquitin chain topology in plant cell signaling: a new facet to an evergreen story. Front Plant Sci. 5, 122.

10. Welchman, R.L., C. Gordon, and R.J. Mayer. (2005) Ubiquitin and ubiquitin-like proteins as multifunctional signals. Nat Rev Mol Cell Biol. 6(8), 599–609.

11. Hershko, A. and A. Ciechanover. (1998) The ubiquitin system. Annu Rev Biochem. 67(1), 425–479.

12. Farrell, P.J., R.J. Broeze, and P. Lengyel. (1979) Accumulation of an mRNA and protein in interferon-treated Ehrlich ascites tumour cells. Nature. 279(5713), 523–525.

13. Kumar, S., Y. Tomooka, and M. Noda. (1992) Identification of a set of genes with developmentally down-regulated expression in the mouse brain. Biochem Biophys Res Commun. 185(3), 1155–1161.

14. Meluh, P.B. and D. Koshland. (1995) Evidence that the MIF2 gene of Saccharomyces cerevisiae encodes a centromere protein with homology to the mammalian centromere protein CENP-C. Mol Biol Cell. 6(7), 793–807.

15. Boddy, M.N., K. Howe, L.D. Etkin, E. Solomon, and P.S. Freemont. (1996) PIC 1, a novel ubiquitin-like protein which interacts with the PML component of a multiprotein complex that is disrupted in acute promyelocytic leukaemia. Oncogene. 13(5), 971–982.

16. Mizushima, N., T. Noda, T. Yoshimori, Y. Tanaka, T. Ishii, et al. (1998) A protein conjugation system essential for autophagy. Nature. 395(6700), 395–398.

17. Furukawa, K., N. Mizushima, T. Noda, and Y. Ohsumi. (2000) A protein conjugation system in yeast with homology to biosynthetic enzyme reaction of prokaryotes. J Biol Chem. 275(11), 7462–7465.

18. 18. Van der Veen, A.G., K. Schorpp, C. Schlieker, L. Buti, J.R. Damon, et al. (2011) Role of the ubiquitin-like protein Urm1 as a noncanonical lysine-directed protein modifier. Proc Natl Acad Sci U S A. 108(5), 1763–1770.

19. Petroski, M.D., G.S. Salvesen, and D.A. Wolf. (2011) Urm1 couples sulfur transfer to ubiquitin-like protein function in oxidative stress. Proc Natl Acad Sci U S A. 108(5), 1749–1750.

20. Noda, N.N., Y. Ohsumi, and F. Inagaki. (2011) Crystallographic Studies on Autophagy-Related Proteins. Current Trends in X-Ray Crystallography. 333.

21. Radoshevich, L., L. Murrow, N. Chen, E. Fernandez, S. Roy, et al. (2010) ATG12 conjugation to ATG3 regulates mitochondrial homeostasis and cell death. Cell. 142(4), 590–600.

22. Gong, P., A. Canaan, B. Wang, J. Leventhal, A. Snyder, et al. (2010) *The ubiquitin-like protein FAT10 mediates NF-*κ*B activation*. J Am Soc Nephrol. 21(2), 316–326.

23. Geng, J. and D. Klionsky. (2008) *The Atg8 and Atg12 ubiquitin*L*like conjugation systems in macroautophagy*. EMBO Rep. 9(9), 859–864.

24. Tanaka, K., J. Nishide, K. Okazaki, H. Kato, O. Niwa, et al. (1999) Characterization of a fission yeast SUMO-1 homologue, pmt3p, required for multiple nuclear events, including the control of telomere length and chromosome segregation. Mol Cell Biol. 19(12), 8660–8672.

25. Cox, J. and M. Mann. (2008) MaxQuant enables high peptide identification rates, individualized ppb-range mass accuracies and proteome-wide protein quantification. Nat Biotechnol. 26(12), 1367–1372.

26. Perkins, D.N., D.J. Pappin, D.M. Creasy, and J.S. Cottrell. (1999) *Probability*L*based protein identification by searching sequence databases using mass spectrometry data*. ELECTROPHORESIS. 20(18), 3551–3567.

27. Eng, J.K., A.L. McCormack, and J.R. Yates. (1994) An approach to correlate tandem mass spectral data of peptides with amino acid sequences in a protein database. J Am Soc Mass Spectrom. 5(11), 976–989.

28. Hendriks, I.A., D. Lyon, D. Su, N.H. Skotte, J.A. Daniel, et al. (2018) Site-specific characterization of endogenous SUMOylation across species and organs. Nat Commun. 9(1), 2456.

29. Lamoliatte, F., F.P. McManus, G. Maarifi, M.K. Chelbi-Alix, and P. Thibault. (2017) Uncovering the SUMOylation and ubiquitylation crosstalk in human cells using sequential peptide immunopurification. Nat Commun. 8(1), 14109.

30. Hendriks, I.A., D. Lyon, C. Young, L.J. Jensen, A.C. Vertegaal, et al. (2017) Site-specific mapping of the human SUMO proteome reveals co-modification with phosphorylation. Nat Struct Mol Biol. 24(3), 325–336.

31. Cai, L., J. Tu, L. Song, Z. Gao, K. Li, et al. (2017) Proteome-wide Mapping of Endogenous SUMOylation Sites in Mouse Testis. Mol Cell Proteomics. 16(5), 717–727.

32. Køhler, J.B., T. Tammsalu, M.M. Jørgensen, N. Steen, R.T. Hay, et al. (2015) Targeting of SUMO substrates to a Cdc48–Ufd1–Npl4 segregase and STUbL pathway in fission yeast. Nat Commun. 6(1), 1–14.

33. Lamoliatte, F., D. Caron, C. Durette, L. Mahrouche, M.A. Maroui, et al. (2014) Large-scale analysis of lysine SUMOylation by SUMO remnant immunoaffinity profiling. Nat Commun. 5(1), 5409.

34. Lobato-Gil, S., J.B. Heidelberger, C. Maghames, A. Bailly, L. Brunello, et al. (2021) Proteome-wide identification of NEDD8 modification sites reveals distinct proteomes for canonical and atypical NEDDylation. Cell Rep. 34(3), 108635.

35. Kim, W., E.J. Bennett, E.L. Huttlin, A. Guo, J. Li, et al. (2011) Systematic and quantitative assessment of the ubiquitin-modified proteome. Mol Cell. 44(2), 325–340.

36. Xu, G., J.S. Paige, and S.R. Jaffrey. (2010) Global analysis of lysine ubiquitination by ubiquitin remnant immunoaffinity profiling. Nat Biotechnol. 28(8), 868–873.

37. Chen, Z.L., J.M. Meng, Y. Cao, J.L. Yin, R.Q. Fang, et al. (2019) A high-speed search engine pLink 2 with systematic evaluation for proteome-scale identification of cross-linked peptides. Nat Commun. 10(1), 3404.

38. Yang, B., Y.J. Wu, M. Zhu, S.B. Fan, J. Lin, et al. (2012) Identification of cross-linked peptides from complex samples. Nat Methods. 9(9), 904–906.

39. Hsiao, H.-H., E. Meulmeester, B.T. Frank, F. Melchior, and H. Urlaub. (2009) *“*ChopNSpice,” a mass spectrometric approach that allows identification of endogenous small ubiquitin-like modifier-conjugated peptides. Mol Cell Proteomics. 8(12), 2664–2675.

40. Sakamaki, J.I. and N. Mizushima. (2023) Ubiquitination of non-protein substrates. Trends Cell Biol. 33(11), 991–1003.

41. Kelsall, I.R. (2022) Non-lysine ubiquitylation: Doing things differently. Front Mol Biosci. 9, 1008175.

42. Ichimura, Y., T. Kirisako, T. Takao, Y. Satomi, Y. Shimonishi, et al. (2000) A ubiquitin-like system mediates protein lipidation. Nature. 408(6811), 488–492.

43. Hanada, T., N.N. Noda, Y. Satomi, Y. Ichimura, Y. Fujioka, et al. (2007) The Atg12-Atg5 conjugate has a novel E3-like activity for protein lipidation in autophagy. J Biol Chem. 282(52), 37298–37302.

44. Xie, Z., U. Nair, and D.J. Klionsky. (2008) Atg8 controls phagophore expansion during autophagosome formation. Mol Biol Cell. 19(8), 3290–3298.

45. Tanida, I., T. Ueno, and E. Kominami. (2004) LC3 conjugation system in mammalian autophagy. Int J Biochem Cell Biol. 36(12), 2503–2518.

46. Mizushima, N., A. Yamamoto, M. Matsui, T. Yoshimori, and Y. Ohsumi. (2004) In vivo analysis of autophagy in response to nutrient starvation using transgenic mice expressing a fluorescent autophagosome marker. Mol Biol Cell. 15(3), 1101–1111.

47. Sakamaki, J.I., K.L. Ode, Y. Kurikawa, H.R. Ueda, H. Yamamoto, et al. (2022) Ubiquitination of phosphatidylethanolamine in organellar membranes. Mol Cell. 82(19), 3677–3692 e3611.

48. Otten, E.G., E. Werner, A. Crespillo-Casado, K.B. Boyle, V. Dharamdasani, et al. (2021) Ubiquitylation of lipopolysaccharide by RNF213 during bacterial infection. Nature. 594(7861), 111–116.

49. Chi, H., C. Liu, H. Yang, W.F. Zeng, L. Wu, et al. (2018) Comprehensive identification of peptides in tandem mass spectra using an efficient open search engine. Nat Biotechnol. 36(11), 1059–1061.

50. Shao, G., Y. Cao, Z. Chen, C. Liu, S. Li, et al. (2021) How to use open-pFind in deep proteomics data analysis?—A protocol for rigorous identification and quantitation of peptides and proteins from mass spectrometry data. Biophys Rep. 7(3), 207.

51. Taylor, D.L., J.C. Ho, A. Oliver, and F.Z. Watts. (2002) Cell-cycle-dependent localisation of Ulp1, a Schizosaccharomyces pombe Pmt3 (SUMO)-specific protease. J Cell Sci. 115(Pt 6), 1113–1122.

52. Melchior, F., M. Schergaut, and A. Pichler. (2003) SUMO: ligases, isopeptidases and nuclear pores. Trends Biochem Sci. 28(11), 612–618.

53. Wei, Y., L.X. Diao, S. Lu, H.T. Wang, F. Suo, et al. (2017) SUMO-Targeted DNA Translocase Rrp2 Protects the Genome from Top2-Induced DNA Damage. Mol Cell. 66(5), 581–596 e586.

54. Yuan, Z.F.e., C. Liu, H.P. Wang, R.X. Sun, Y. Fu, et al. (2012) *pParse: A method for accurate determination of monoisotopic peaks in high*L*resolution mass spectra*. Proteomics. 12(2), 226–235.

55. Prudden, J., S. Pebernard, G. Raffa, D.A. Slavin, J.J.P. Perry, et al. (2007) *SUMO*L*targeted ubiquitin ligases in genome stability*. The EMBO journal. 26(18), 4089–4101.

56. Pan, Z.Q., A. Kentsis, D.C. Dias, K. Yamoah, and K. Wu. (2004) Nedd8 on cullin: building an expressway to protein destruction. Oncogene. 23(11), 1985–1997.

57. Osaka, F., M. Saeki, S. Katayama, N. Aida, A. Toh-e, et al. (2000) Covalent modifier NEDD8 is essential for SCF ubiquitin-ligase in fission yeast. The EMBO journal. 19(13), 3475.

58. Hofer, S.J., I. Daskalaki, M. Bergmann, J. Friščić, A. Zimmermann, et al. (2024) Spermidine is essential for fasting-mediated autophagy and longevity. Nat Cell Biol. 26(9), 1571–1584.

59. Dever, T.E. and I.P. Ivanov. (2018) Roles of polyamines in translation. J Biol Chem. 293(48), 18719–18729.

60. Minois, N. (2014) Molecular basis of the ‘anti-aging’effect of spermidine and other natural polyamines-a mini-review. Gerontology. 60(4), 319–326.

61. Dever, T.E., E. Gutierrez, and B.S. Shin. (2014) The hypusine-containing translation factor eIF5A. Crit Rev Biochem Mol Biol. 49(5), 413–425.

62. Nie, M., A.A. Vashisht, J.A. Wohlschlegel, and M.N. Boddy. (2015) High Confidence Fission Yeast SUMO Conjugates Identified by Tandem Denaturing Affinity Purification. Sci Rep. 5(1), 14389.

63. Sagar, N.A., S. Tarafdar, S. Agarwal, A. Tarafdar, and S. Sharma. (2021) Polyamines: Functions, Metabolism, and Role in Human Disease Management. Med Sci (Basel). 9(2).

64. Michael, A.J. (2016) Biosynthesis of polyamines and polyamine-containing molecules. Biochem J. 473(15), 2315–2329.

65. Igarashi, K. and K. Kashiwagi. (2015) Modulation of protein synthesis by polyamines. IUBMB Life. 67(3), 160–169.

66. Chattopadhyay, M.K., M.H. Park, and H. Tabor. (2008) Hypusine modification for growth is the major function of spermidine in Saccharomyces cerevisiae polyamine auxotrophs grown in limiting spermidine. Proc Natl Acad Sci U S A. 105(18), 6554–6559.

67. Pederick, J.L., J. Klose, B. Jovcevski, T.L. Pukala, and J.B. Bruning. (2023) *Escherichia coli YgiC and YjfC Possess Peptide*─ *Spermidine Ligase Activity*. Biochemistry. 62(4), 899–911.

68. Singh, V.P., S. Hirose, M. Takemoto, A. Farrag, S.I. Sato, et al. (2024) Chemoproteomic Identification of Spermidine-Binding Proteins and Antitumor-Immunity Activators. J Am Chem Soc.

69. O’Shea, J.P., M.F. Chou, S.A. Quader, J.K. Ryan, G.M. Church, et al. (2013) pLogo: a probabilistic approach to visualizing sequence motifs. Nat Methods. 10(12), 1211–1212.

70. Ma, J., T. Chen, S. Wu, C. Yang, M. Bai, et al. (2019) iProX: an integrated proteome resource. Nucleic Acids Res. 47(D1), D1211–D1217.

71. Chen, T., J. Ma, Y. Liu, Z. Chen, N. Xiao, et al. (2022) iProX in 2021: connecting proteomics data sharing with big data. Nucleic Acids Res. 50(D1), D1522–D1527.

