## Supplemental Figures for "Global analysis of protein and small-molecule substrates of ubiquitin-like proteins (UBLs)"

Shao et al.

### Supplementary Note

#### Supplementary Note 1. Selection of ion types in the fine-scoring algorithm of pLink-UBL

To analyze the fragmentation characteristics of SUMOylated peptides and further optimize pLink-UBL, we synthesized two SUMO peptides, Pmt3 and SUMO2, and 246 substrate peptides from 30 peptide templates **(Supplementary Fig. S4).** Each substrate peptide template was chemically linked to each of the two SUMO peptides, resulting in a total of 492 SUMO-substrate peptide pairs. After LC-MS/MS analysis, a total of 1,970 Pmt3 SUMOylated spectra and a total of 1,560 SUMO2 SUMOylated spectra were manually annotated. These spectral collections were named the synthetic-Pmt3 dataset and the synthetic-SUMO2 dataset, respectively.

After initial data exploration, we found that the sequence coverages of SUMO peptides were higher than those of substrate peptides, suggestive of better fragmentation of the SUMO peptides than substrate peptides **(Supplementary Fig. S2)**. We therefore developed a two-stage search strategy in pLink-UBL. It first identifies the SUMO peptide, then the substrate peptide. To further optimize the fine-scoring algorithm of pLink-UBL, significance of matching of each type of theoretical fragment ions with experimental peaks was analyzed **(Supplementary Fig. S5 and S6)**, and top 18 fragment ion types were selected for fine scoring.

#### Supplementary Note 2. pLink-UBL identifies UBL-C-pep before it identifies the cross-link between a UBL-C-pep and a substrate peptide

In the first stage, all possible SUMO peptides are generated from *in silico* enzymatic digestion of the SUMO protein according to a user's choice of enzyme and the maximum missed cleavage sites. Then the SUMO peptides thus generated are coarse-scored with each spectrum as described previously [1]. Only the top-3 coarse-scored SUMO peptide candidates are kept. As SUMO peptides are easier to identify than substrate peptides, only *b/y* ions of charge state 1+ or 2+ of SUMO peptides are used for coarse-scoring. In the second stage, the mass of the candidate substrate peptide, equal to the precursor mass minus the mass of the candidate SUMO peptide, is used to retrieve the sequence of a candidate substrate peptide from a peptide index. As previously described [1], the peptide index is a hash table, which maps a mass to all peptides within a user-defined mass tolerance. With the peptide index, pLink-UBL retrieves the candidate substrate peptide at a fixed time cost regardless of the size of the protein database. Once the candidate SUMO peptide and the candidate substrate peptides are retrieved, they are linked *in silico* and fine-scored against the MS2 spectrum. The fine-scoring algorithm is the same as that in pLink 2 [1], except that fragment ions used for fine-scoring are optimized for SUMOylated peptides **(Supplementary Note 1, and Supplementary Fig. S5 and S6)**. Only the top-ranked result is kept for each spectrum after fine scoring.

### Supplemental Figures

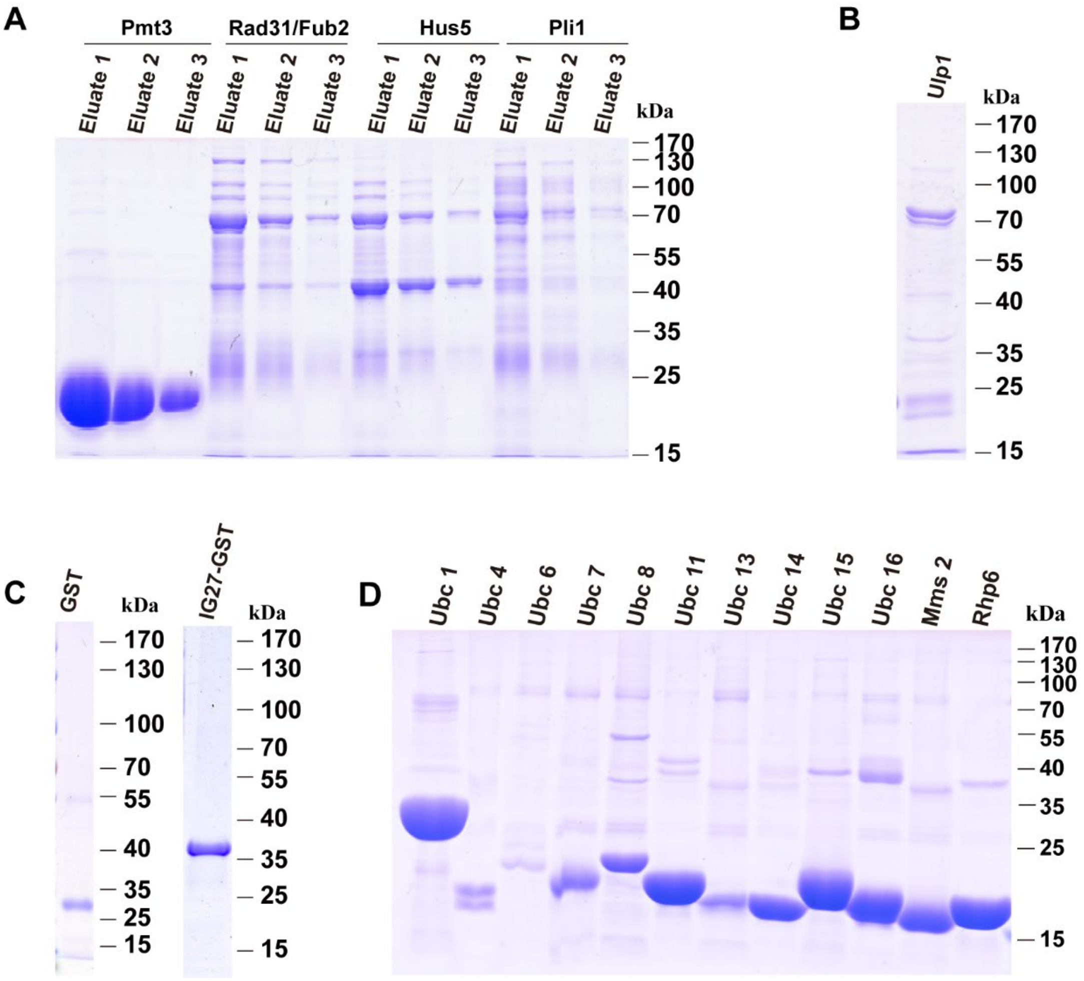


**Supplementary Figure S1. Protein expression in *E. coli.* (A)** Expression of Pmt3, Rad31/Fub2, Hus5 and pli1. **(B)** Expression of Ulp1. **(C)** Expression of GST and IG27-GST. **(D)** Expression of Ubc1, Ubc4, Ubc6, Ubc7, Ubc11, Ubc13, Ubc14, Ubc15, Ubc16, Mms2 and Rhp6.

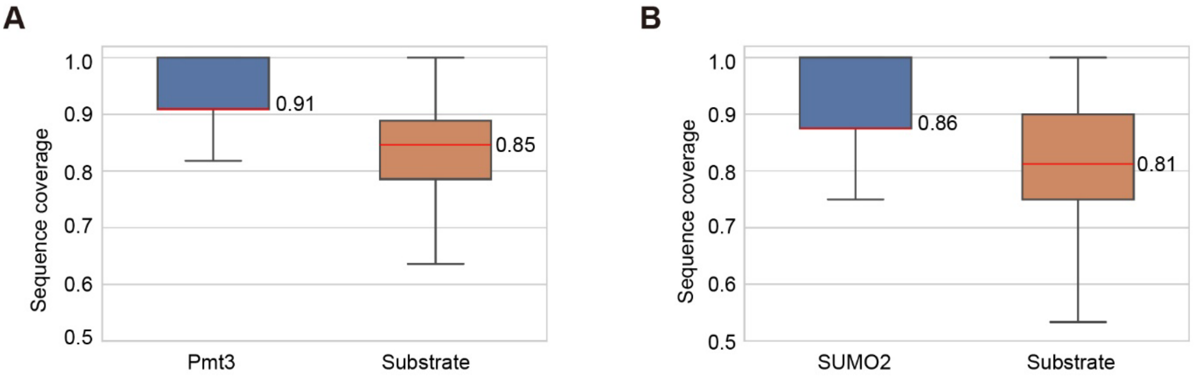


**C**

**Synthetic-Pmt3**

| Ion type | $Prob_{occur}$ | Average intensity |
|---|---|---|
| $b^{1+}$ | 0.59 | 0.12 |
| $b^{2+}$ | 0.16 | 0.03 |
| $y_{1+}$ | 0.62 | 0.17 |
| $y^{2+}$ | 0.38 | 0.07 |

**D**

**Synthetic-SUMO2**

| Ion type | $Prob_{occur}$ | Average intensity |
|---|---|---|
| $b^{1+}$ | 0.55 | 0.09 |
| $b^{2+}$ | 0.17 | 0.04 |
| $y_{1+}$ | 0.63 | 0.21 |
| $y^{2+}$ | 0.35 | 0.06 |

**Supplementary Figure S2. Fragmentation characteristics of synthetic SUMOylated peptides in Pmt3 and SUMO2 libraries.** (**A)** Distribution of sequence coverage for the Synthetic-Pmt3 dataset. **(B)** Distribution of sequence coverage for the Synthetic-SUMO2 dataset. The sequence coverage was calculated as the number of cleavage sites supported by matching $b^{1+}/b^{2+}/y^{1+}/y^{2+}$ peaks divided by the length at the sequence. Box indicates the interquartile range (IQR) and its whiskers 1.5×IQR values; no outliers are shown. The median sequence coverage values are indicated. For both Pmt3 and SUMO2 peptides, the median equals to the lower quartile, and the upper whisker equals to the upper quartile. (**C)** Occurrence probabilities and average intensities of ion types $b^{1+}$, $b^{2+}$, $y^{1+}$, and $y^{2+}$ for the Synthetic-Pmt3 dataset. The occurrence probability is the number of matched peaks of an ion type divided by the total number of theoretical peaks of that ion type. The average intensity is the average of the normalized intensity of matched peaks of the indicated ion type (normalized to the base peak intensity). (**D)** Similar to (C), but for the Synthetic-SUMO2 dataset.

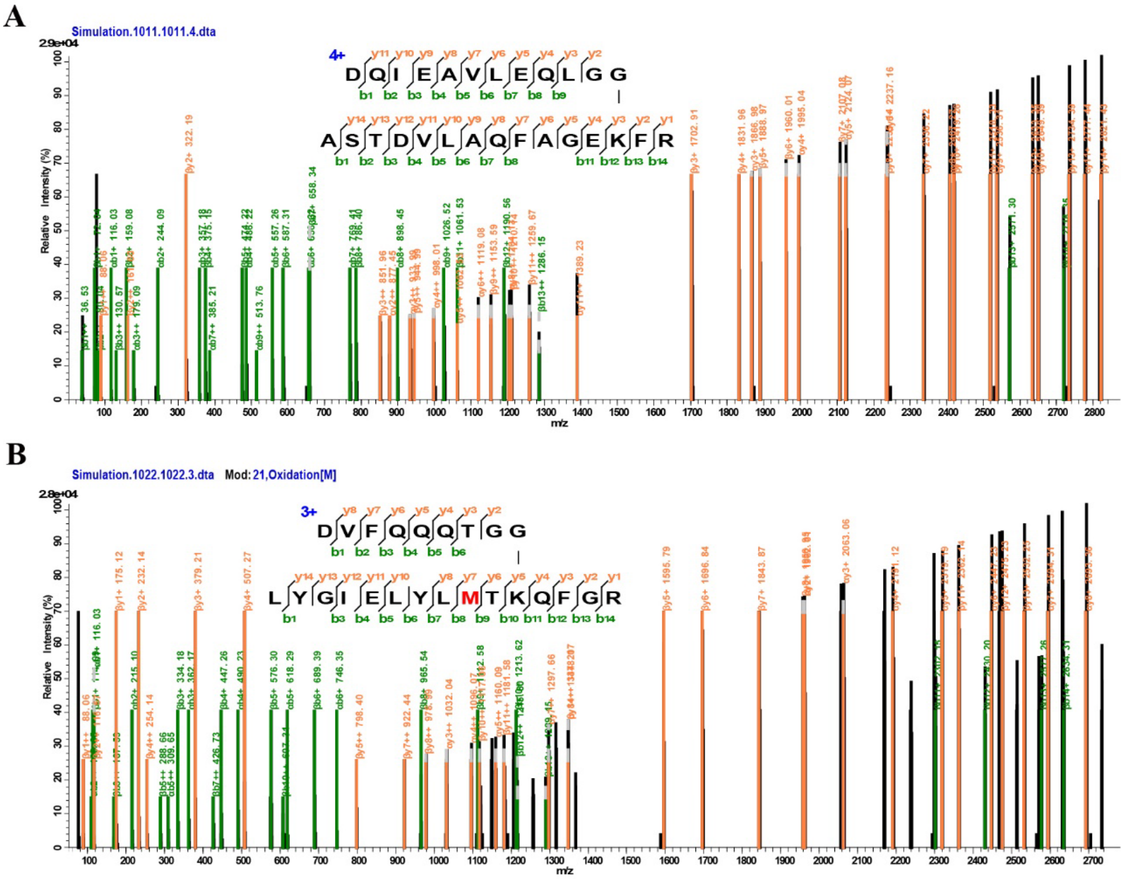


**Supplementary Figure S3.** Two examples of simulated spectra SUMOylated by **(A)** Pmt3 and **(B)** SUMO2, respectively, and annotated by pLabel [2].

| Name | $H_2N$—DQIEAVLEQLGG—C(=O)—lysine (K) of mod_pep_X $H_2N$—DVFQQQTGG—C(=O)—lysine (K) of mod_pep_X<br>mod_pep_x | Peptide count |
|---|---|---|
| **SP1** | D(V/L/A)KPSTEHI(H/W)LK(3) | 6 |
| **SP2** | VK(V/G/S)YSPS(E/F)EK(2) | 6 |
| **SP3** | NSLP(N/L/F/Y/A/G)PVKIQR(8) | 6 |
| **SP4** | AT(E/G/W)VK(A/S/T)DFRAR(5) | 9 |
| **SP5** | E(F/W/V)KAESLA(L/S/T)IK(3) | 9 |
| **SP6** | TALH(G/T/L)WS(Y/A/H)TKFR(10) | 9 |
| **SP7** | DL(Q/F/T)LHTK(N/G/W)EHLEK(7) | 9 |
| **SP8** | KLQP(S/L/R)SKPLSSL(E/D/F)AR(7) | 9 |
| **SP9** | D(S/N/F)IKTPHT(E/D)LQK(4) | 6 |
| **SP10** | EAKHEDD(A/W)NT(L/T/H)QPK | 6 |
| **SP11** | L(D/E)AELKLAGEY(G/A/V)LR(6) | 6 |
| **SP12** | VFE(N/S/T)VKQEN(P/H)DEK(6) | 6 |
| **SP13** | ARPQ(V/L/Y)VPKAA(E/D)PK(8) | 6 |
| **SP14** | (Y/L/A)SKAYEAYQQA(D/E/G)YR(3) | 9 |
| **SP15** | AT(A/S/T)WKP(E/G/W)FRAK(5) | 9 |
| **SP16** | NTE(N/G/Y)STKEEGTWE(V/L/S)NK(7) | 9 |
| **SP17** | D(P/G/W)NLFKFEGAGT(S/N/H)LR(6) | 9 |
| **SP18** | VV(G/T/H)QDNNEV(A/Y/H)FKIK(12) | 9 |
| **SP19** | AYSATIP(G/L/Y)KNES(S/V/E)AK(9) | 9 |
| **SP20** | LQTS(E/D/G)SLEKNH(L/A/Y)K(9) | 9 |
| **SP21** | AGS(D/E/F)KIVSH(T/G/Y)LSGHK(5) | 9 |
| **SP22** | DI(Y/L/G)KVAFTGS(D/E/F)GVGR(4) | 9 |
| **SP23** | D(S/V/N)LKLEECIES(W/A/E)SLNQK(4) | 9 |
| **SP24** | DS(A/G/E)VKQEGNLLIHP(L/H/Y)NSLK(5) | 9 |
| **SP25** | DMKEYL(E/D/G)AGLVG(L/A/Y)EK(3) | 9 |
| **SP26** | VDSKSPIPF(S/N/T)VP(P/H/G)SR(4) | 9 |
| **SP27** | TADTDVS(G/A/S/V/L/E/H/F/Y)TEPVKR(13) | 9 |
| **SP28** | EMALIPKSEEDL(G/A/S/P/L/E/H/F/Y)NER(7) | 9 |
| **SP29** | KDE(F/A/G)STQKIQQ(Y/H/L)VVR(8) | 9 |
| **SP30** | EVSS(L/A/Y)ESQLKLAN(E/D/G)R(10) | 9 |

**Supplementary Figure S4. The list of synthetic SUMOylated peptides.** The SUMOylated lysine residue is red. In parentheses at given are possible amino acids at the indicated position. Peptide count is the number of all possible SUMOylated peptides in the sample.

| # | Ion Type | Peak Ratio | Ion Ratio | Avg Int | Sig | # Spec containing this ion type | # Peaks / Spec |
|---|---|---|---|---|---|---|---|
| 1 | $b^{1+}$ | 9.74E-02 | 7.55E-01 | 2.07E-01 | 1.52E-02 | 1,970 | 12.17 |
| 2 | $y^{1+}$ | 5.71E-02 | 8.67E-01 | 2.18E-01 | 1.08E-02 | 1,970 | 7.27 |
| 3 | $yb^{1+}$ | 2.36E-01 | 4.49E-01 | 7.52E-02 | 7.96E-03 | 1,970 | 29.92 |
| 4 | $y^{2+x}$ | 6.12E-02 | 5.26E-01 | 8.46E-02 | 2.72E-03 | 1,943 | 8.21 |
| 5 | $b^{1+}$-$H_2O$ | 7.43E-02 | 5.96E-01 | 4.33E-02 | 1.92E-03 | 1,970 | 9.59 |
| 6 | $y^{1+x}$ | 4.23E-02 | 3.69E-01 | 1.16E-01 | 1.80E-03 | 1,652 | 5.82 |
| 7 | $b^{1+}$-$NH_3$ | 6.06E-02 | 4.97E-01 | 5.62E-02 | 1.69E-03 | 1,970 | 7.87 |
| 8 | $a^{1+}$ | 5.86E-02 | 4.80E-01 | 4.63E-02 | 1.30E-03 | 1,970 | 7.65 |
| 9 | $p^{4+x}$ | 9.70E-03 | 5.11E-01 | 1.99E-01 | 9.85E-04 | 525 | 1.02 |
| 10 | $ya^{1+}$ | 1.03E-01 | 2.02E-01 | 3.69E-02 | 7.66E-04 | 1,970 | 13.25 |
| 11 | $y^{2+x}$-$NH_3$ | 4.38E-02 | 3.86E-01 | 4.35E-02 | 7.35E-04 | 1,900 | 6.05 |
| 12 | $y^{1+}$-$H_2O$ | 2.90E-02 | 4.55E-01 | 3.95E-02 | 5.21E-04 | 1,801 | 3.90 |
| 13 | $y^{1+x}$-$NH_3$ | 3.27E-02 | 3.03E-01 | 4.36E-02 | 4.32E-04 | 1,334 | 4.79 |
| 14 | $yb^{1+x}$ | 4.21E-02 | 3.12E-01 | 3.13E-02 | 4.11E-04 | 1,828 | 5.90 |
| 15 | $v^{1+}$ | 2.51E-02 | 2.22E-01 | 7.12E-02 | 3.97E-04 | 1,752 | 3.51 |
| 16 | $y^{1+}$-$NH_3$ | 2.59E-02 | 4.25E-01 | 3.36E-02 | 3.70E-04 | 1,800 | 3.60 |
| 17 | $p^{1+}$ | 8.38E-03 | 5.80E-01 | 7.48E-02 | 3.63E-04 | 1,344 | 1.16 |
| 18 | $y^{2+x}$-$H_2O$ | 3.70E-02 | 3.31E-01 | 2.79E-02 | 3.41E-04 | 1,812 | 5.16 |
| 19 | $b^{1+x}$ | 2.37E-02 | 4.23E-01 | 3.29E-02 | 3.31E-04 | 612 | 3.83 |
| 20 | $y^{1+x}$-$H_2O$ | 2.66E-02 | 2.54E-01 | 3.26E-02 | 2.20E-04 | 1,144 | 3.96 |
| 21 | $y^{2+}$ | 1.78E-02 | 2.42E-01 | 5.08E-02 | 2.19E-04 | 722 | 2.33 |
| 22 | $p^{1+x}$ | 9.47E-03 | 7.14E-01 | 2.39E-02 | 1.62E-04 | 983 | 1.43 |
| 23 | $p^{2+}$ | 7.99E-03 | 5.10E-01 | 3.45E-02 | 1.41E-04 | 1,116 | 1.02 |
| 24 | $u^{1+}$ | 1.86E-02 | 3.21E-01 | 2.27E-02 | 1.35E-04 | 1,471 | 2.76 |
| 25 | $b^{2+x}$ | 1.41E-02 | 2.42E-01 | 3.44E-02 | 1.17E-04 | 1,106 | 2.12 |
| 26 | $y^{3+x}$-$NH_3$ | 1.73E-02 | 1.41E-01 | 3.80E-02 | 9.24E-05 | 853 | 2.23 |
| 27 | $y^{3+x}$ | 1.77E-02 | 1.43E-01 | 3.50E-02 | 8.85E-05 | 734 | 2.28 |
| 28 | $b^{2+x}$-$NH_3$ | 1.30E-02 | 2.01E-01 | 3.25E-02 | 8.49E-05 | 901 | 1.93 |
| 29 | $b^{2+x}$-$H_2O$ | 1.26E-02 | 1.98E-01 | 3.31E-02 | 8.26E-05 | 984 | 1.90 |
| 30 | $a^{2+x}$ | 1.21E-02 | 2.04E-01 | 2.83E-02 | 6.95E-05 | 1,094 | 1.78 |
| 31 | $y^{3+x}$-$H_2O$ | 1.49E-02 | 1.21E-01 | 3.56E-02 | 6.43E-05 | 801 | 1.89 |
| 32 | $v^{2+}$ | 1.49E-02 | 1.25E-01 | 3.38E-02 | 6.26E-05 | 1,151 | 1.93 |
| 33 | $ya^{1+x}$ | 1.65E-02 | 1.27E-01 | 2.94E-02 | 6.15E-05 | 1,168 | 2.41 |
| 34 | $ya^{2+x}$ | 1.27E-02 | 8.97E-02 | 4.87E-02 | 5.55E-05 | 1,330 | 1.69 |
| 35 | $y^{2+}$-$H_2O$ | 1.11E-02 | 1.49E-01 | 3.29E-02 | 5.44E-05 | 512 | 1.46 |
| 36 | $yb^{3+x}$ | 1.30E-02 | 8.36E-02 | 4.76E-02 | 5.16E-05 | 557 | 1.59 |
| 37 | $y^{2+}$-$NH_3$ | 9.45E-03 | 1.28E-01 | 4.13E-02 | 5.01E-05 | 536 | 1.23 |
| 38 | $b^{3+}$ | 9.01E-03 | 6.56E-02 | 8.33E-02 | 4.93E-05 | 1,296 | 1.03 |
| 39 | $yb^{2+x}$ | 1.43E-02 | 1.02E-01 | 3.31E-02 | 4.83E-05 | 1,361 | 1.94 |
| 40 | $u^{2+}$ | 9.61E-03 | 1.49E-01 | 2.20E-02 | 3.15E-05 | 535 | 1.35 |
| 41 | $b^{2+}$ | 1.17E-02 | 8.71E-02 | 2.08E-02 | 2.11E-05 | 671 | 1.41 |
| 42 | $ya^{3+x}$ | 1.12E-02 | 7.38E-02 | 2.48E-02 | 2.05E-05 | 505 | 1.43 |
| 43 | $b^{2+}$-$NH_3$ | 8.39E-03 | 6.70E-02 | 2.95E-02 | 1.66E-05 | 548 | 1.06 |

**Supplementary Figure S5. The list of ion types ranked by significance of matching on the Synthetic-Pmt3 dataset.** The peak ratio is the number of matched peaks of an ion type divided by the total number of matched peaks. The ion ratio is the number of matched peaks of an ion type divided by the total number of theoretical peaks of that ion type. The average intensity (Avg Int) is the average intensity of all matched peaks normalized against the base peak. The matching significance (Sig) is the product of the peak ratio, ion ratio, and average intensity.

| # | Ion Type | Peak Ratio | Ion Ratio | Avg Int | Sig | # Spec containing this ion type | # Peaks / Spec |
|---|---|---|---|---|---|---|---|
| 1 | $y^{1+}$ | 7.17E-02 | 8.78E-01 | 2.85E-01 | 1.79E-02 | 1,560 | 6.65 |
| 2 | $b^{1+}$ | 9.78E-02 | 6.72E-01 | 1.33E-01 | 8.77E-03 | 1,560 | 9.08 |
| 3 | $yb^{1+}$ | 1.98E-01 | 3.51E-01 | 9.34E-02 | 6.48E-03 | 1,560 | 19.01 |
| 4 | $y^{1+x}$ | 4.97E-02 | 3.81E-01 | 1.37E-01 | 2.59E-03 | 1,169 | 5.12 |
| 5 | $a^{1+}$ | 3.88E-02 | 2.93E-01 | 2.16E-01 | 2.46E-03 | 1,532 | 3.89 |
| 6 | $v^{1+}$ | 5.08E-02 | 3.60E-01 | 1.09E-01 | 1.99E-03 | 1,560 | 4.75 |
| 7 | $y^{2+x}$ | 5.60E-02 | 4.38E-01 | 6.41E-02 | 1.57E-03 | 1,483 | 5.69 |
| 8 | $y^{1+x}$-$NH_3$ | 4.65E-02 | 3.91E-01 | 6.49E-02 | 1.18E-03 | 820 | 5.21 |
| 9 | $y^{2+x}$-$NH_3$ | 4.85E-02 | 3.92E-01 | 5.90E-02 | 1.12E-03 | 1,479 | 5.08 |
| 10 | $p^{2+}$ | 1.54E-02 | 7.19E-01 | 1.00E-01 | 1.11E-03 | 1,555 | 1.44 |
| 11 | $b^{1+}$-$H_2O$ | 5.94E-02 | 4.44E-01 | 3.76E-02 | 9.92E-04 | 1,533 | 6.06 |
| 12 | $y^{1+}$-$H_2O$ | 3.66E-02 | 4.63E-01 | 5.10E-02 | 8.64E-04 | 1,402 | 3.68 |
| 13 | $y^{1+}$-$NH_3$ | 3.62E-02 | 4.75E-01 | 4.29E-02 | 7.37E-04 | 1,249 | 3.75 |
| 14 | $b^{1+x}$ | 2.97E-02 | 4.27E-01 | 4.76E-02 | 6.04E-04 | 569 | 3.56 |
| 15 | $y^{2+x}$-$H_2O$ | 4.06E-02 | 3.35E-01 | 4.26E-02 | 5.78E-04 | 1,388 | 4.31 |
| 16 | $y^{1+x}$-$H_2O$ | 3.53E-02 | 3.07E-01 | 4.89E-02 | 5.30E-04 | 708 | 4.03 |
| 17 | $yb^{1+x}$ | 5.03E-02 | 2.99E-01 | 3.42E-02 | 5.13E-04 | 1,390 | 5.59 |
| 18 | $p^{1+}$ | 9.82E-03 | 5.16E-01 | 9.65E-02 | 4.89E-04 | 785 | 1.03 |
| 19 | $ya^{1+}$ | 7.44E-02 | 1.41E-01 | 4.15E-02 | 4.34E-04 | 1,507 | 7.46 |
| 20 | $b^{1+}$-$NH_3$ | 4.03E-02 | 3.21E-01 | 3.23E-02 | 4.17E-04 | 1,424 | 4.25 |
| 21 | $p^{1+x}$ | 1.20E-02 | 6.40E-01 | 5.37E-02 | 4.14E-04 | 1,008 | 1.28 |
| 22 | $y^{2+}$ | 2.16E-02 | 2.52E-01 | 6.08E-02 | 3.31E-04 | 582 | 2.09 |
| 23 | $y^{3+x}$-$NH_3$ | 2.62E-02 | 1.94E-01 | 3.93E-02 | 2.00E-04 | 614 | 2.49 |
| 24 | $u^{1+}$ | 2.03E-02 | 2.87E-01 | 3.16E-02 | 1.84E-04 | 970 | 2.31 |
| 25 | $b^{1+x}$-$NH_3$ | 1.92E-02 | 2.86E-01 | 3.18E-02 | 1.75E-04 | 464 | 2.44 |
| 26 | $b^{2+x}$ | 1.76E-02 | 2.54E-01 | 3.20E-02 | 1.43E-04 | 746 | 2.05 |
| 27 | $y^{3+x}$ | 2.07E-02 | 1.51E-01 | 4.54E-02 | 1.42E-04 | 428 | 1.96 |
| 28 | $a^{1+x}$ | 1.52E-02 | 2.23E-01 | 4.00E-02 | 1.36E-04 | 396 | 1.80 |
| 29 | $y^{3+x}$-$H_2O$ | 2.40E-02 | 1.82E-01 | 2.99E-02 | 1.30E-04 | 577 | 2.34 |
| 30 | $ya^{1+x}$ | 2.37E-02 | 1.47E-01 | 3.69E-02 | 1.29E-04 | 1,030 | 2.71 |
| 31 | $b^{1+x}$-$H_2O$ | 1.54E-02 | 2.24E-01 | 3.42E-02 | 1.18E-04 | 405 | 1.99 |
| 32 | $b^{2+x}$-$NH_3$ | 1.70E-02 | 2.17E-01 | 2.88E-02 | 1.06E-04 | 652 | 1.92 |
| 33 | $a^{2+x}$ | 1.50E-02 | 2.15E-01 | 2.76E-02 | 8.90E-05 | 669 | 1.74 |
| 34 | $b^{2+x}$-$H_2O$ | 1.49E-02 | 2.00E-01 | 2.81E-02 | 8.37E-05 | 713 | 1.73 |
| 35 | $v^{2+}$ | 1.98E-02 | 1.42E-01 | 2.93E-02 | 8.25E-05 | 685 | 1.81 |
| 36 | $yb^{2+}$ | 1.86E-02 | 3.79E-02 | 1.09E-01 | 7.68E-05 | 557 | 1.97 |
| 37 | $b^{3+}$-$H_2O$ | 1.23E-02 | 7.81E-02 | 7.48E-02 | 7.18E-05 | 838 | 1.01 |
| 38 | $yb^{2+x}$ | 1.54E-02 | 8.85E-02 | 2.68E-02 | 3.65E-05 | 759 | 1.64 |
| 39 | $ya^{2+x}$ | 1.44E-02 | 8.01E-02 | 3.04E-02 | 3.52E-05 | 635 | 1.47 |
| 40 | $b^{2+}$-$H_2O$ | 9.10E-03 | 7.73E-02 | 2.31E-02 | 1.62E-05 | 436 | 1.04 |

**Supplementary Figure S6. The list of ion types ranked by significance of matching on the Synthetic-SUMO2 dataset.**

| Charge | Ion type | Notes |
|---|---|---|
| 1~3 | *b* | backbone fragment ions |
| | *y* | |
| | *a* | |
| | *b*-$NH_3$ | Mass of backbone fragment ions - 17.027 Da |
| | *y*-$NH_3$ | |
| | *b*-$H_2O$ | Mass of backbone fragment ions - 18.011 Da |
| | *y*-$H_2O$ | |
| 1~4 | *yb* | The internal ion formed by the two breaks has a *y* ion at one end and a *b* ion at the other end. |
| | *ya* | The internal ion formed by the two breaks has a *y* ion at one end and *a* ion at the other end. |
| | *p* | The mass of a single peptide after cleavage of isopeptide bonds |
| | *u* | *b*-like ions of *p* ion |
| | *v* | *y*-like ions of *p* ion |

**Supplementary Figure S7. Description of ion types**

| Ion types | | |
|---|---|---|
| $b^{1+}$ | $y^{1+}$-$H_2O$ | $yb^{1+}$ |
| $b^{1+}$-$H_2O$ | $y^{1+}$-$NH_3$ | $ya^{1+}$ |
| $b^{1+}$-$NH_3$ | $y^{1+x}$-$H_2O$ | $a^{1+}$ |
| $y^{1+}$ | $y^{1+x}$-$NH_3$ | $p^{1+}$ |
| $y^{1+x}$ | $y^{2+x}$-$H_2O$ | $P^{2+}$ |
| $y^{2+x}$ | $y^{2+x}$-$NH_3$ | $v^{1+}$ |

**Supplementary Figure S8. 18 ion types used in the fine-scoring algorithm of pLink-UBL**

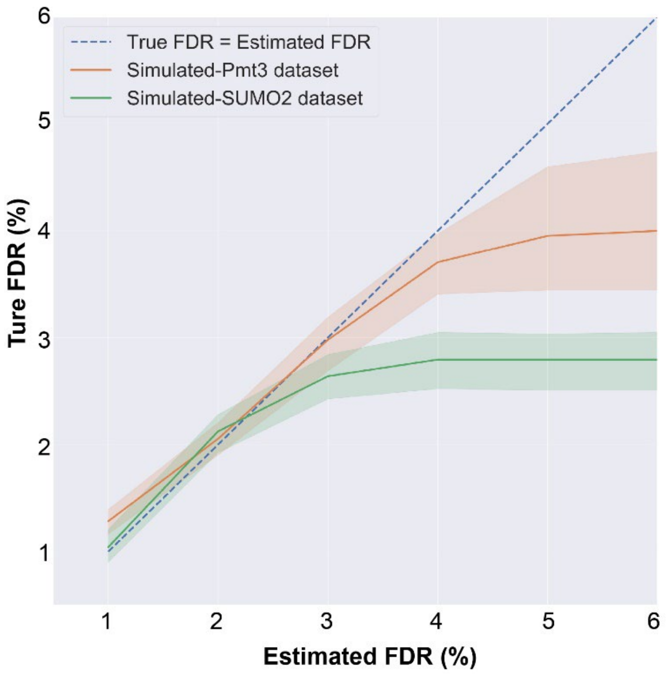


**Supplementary Figure S9. The true FDR compared with the estimated FDR**. No target or decoy PSMs are reported at FDR > 3%. As a result, the curves leveled off.

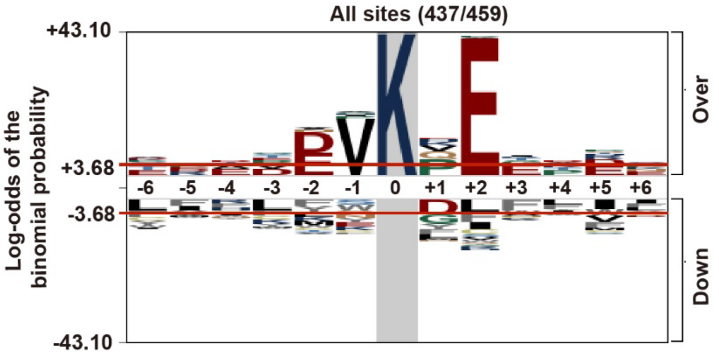


**Supplementary Figure S10. Motif analysis of the sequence surrounding a SUMOylated lysine based on the identification results in this study using pLogo [3].** In parentheses are the number of SUMOylation sites flanked by 6 amino acids on each side, followed by the total number of SUMOylation sites in the category. The log-odds of the binomial probability are presented on the y-axis. Significance threshold values of 3.68 ($p < 0.05$) are visually highlighted in red.

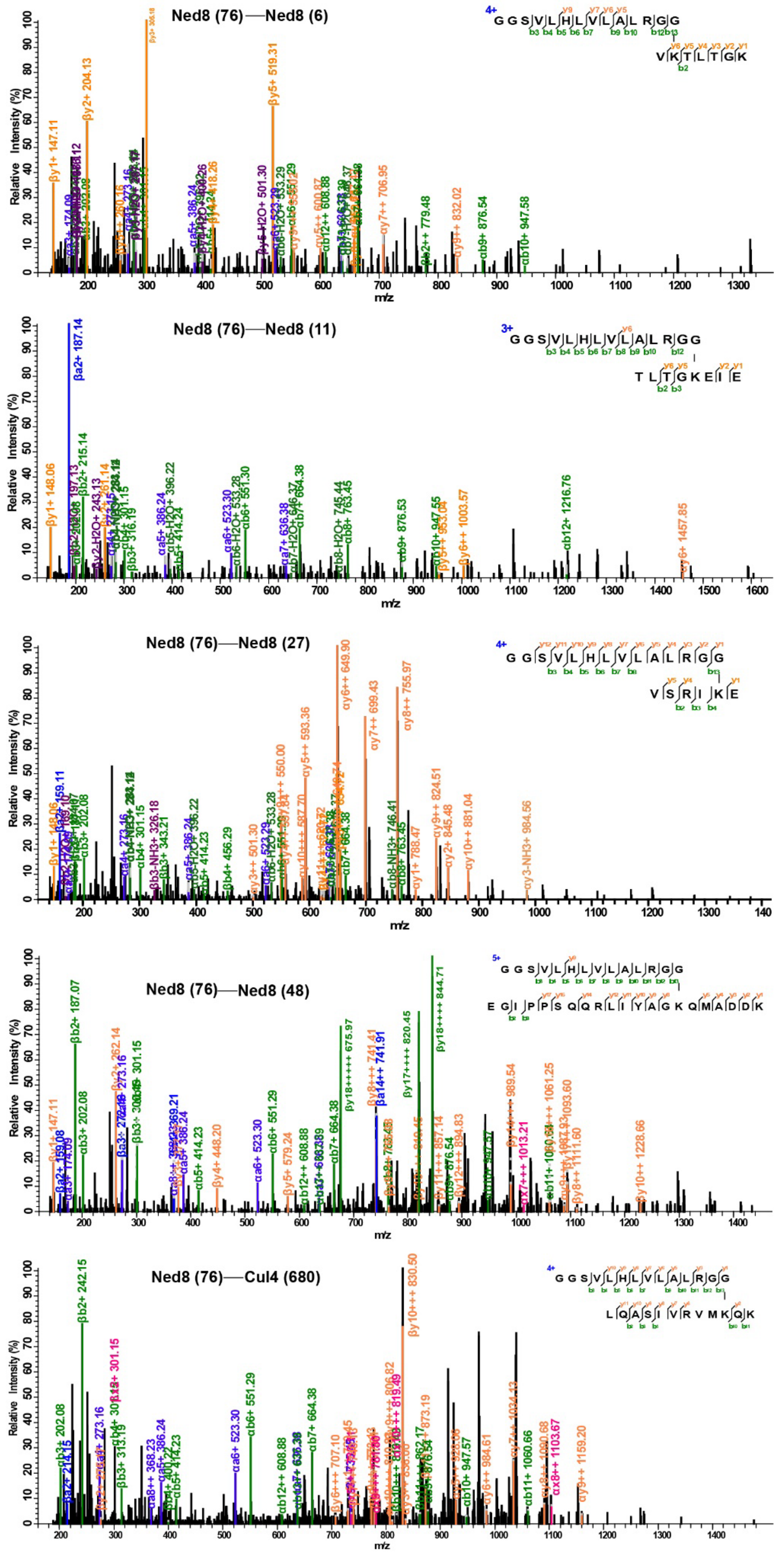


**Supplementary Figure S11. MS2 spectra of five Ned8-modified peptides/sites identified by pLink-UBL.**

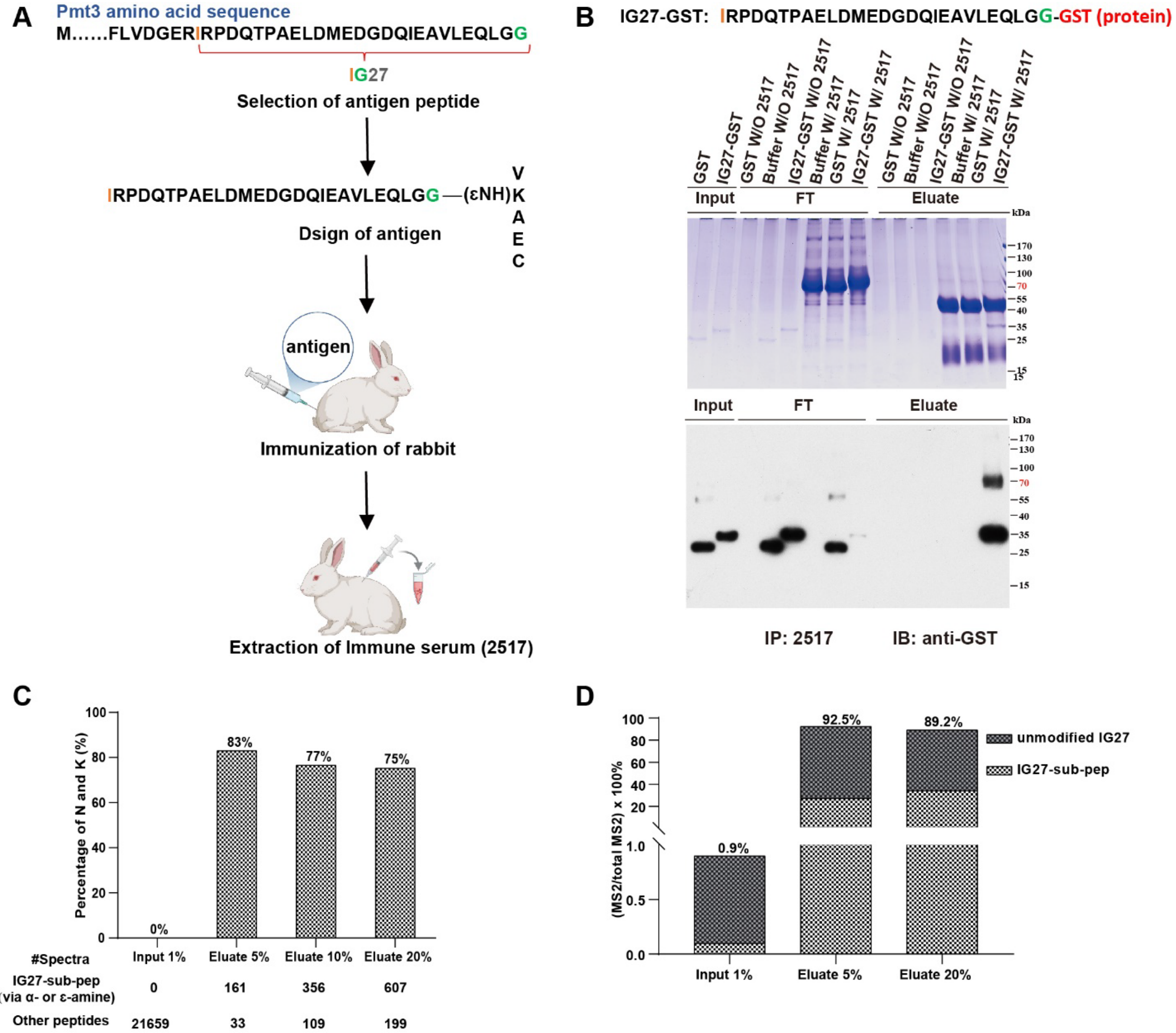


| #Spectra | Input 1% | Eluate 5% | Eluate 10% | Eluate 20% |
|---|---|---|---|---|
| IG27-sub-pep (via α- or ε-amine) | 0 | 161 | 356 | 607 |
| Other peptides | 21659 | 33 | 109 | 199 |

**Supplementary Figure S12. Development and evaluation of an antibody against the C-terminal peptide of Pmt3. (A)** Development of polyclonal antibody 2517 (anti-IG27). **(B)** Evaluation of affinity binding of immune sera using IG27-GST. FT indicates flow through. Sample loading: 1.125% Input, 1.125% FT, 15% Eluate. **(C)** Evaluation of the anti-IG27 antibody using synthetic SUMOylated peptides. **(D)** Evaluation of the anti-IG27 antibody using a sample from fission yeast.

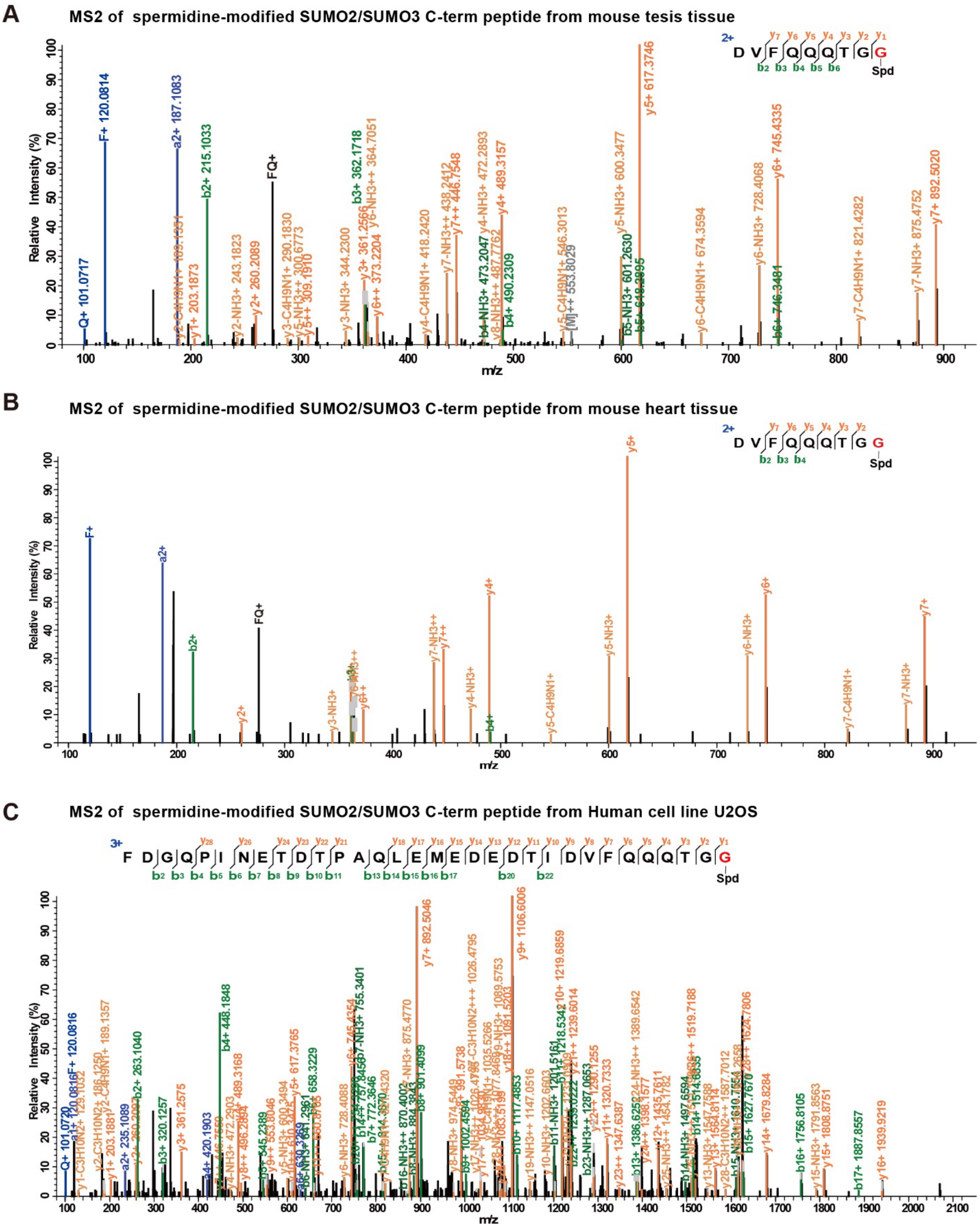


**Supplementary Figure S13. Evidence of SUMO-spermidine conjugation in mouse heart, testis, and human cell line U2OS.** Representative MS2 of mouse (A-B) and human (C) SUMO2/3 C-terminal peptide with spermidine attached at the C terminus.

**SUPPLEMENTARY REFERENCE**


1. Chen, Z.L., J.M. Meng, Y. Cao, J.L. Yin, R.Q. Fang, et al. (2019) *A high-speed search engine pLink 2 with systematic evaluation for proteome-scale identification of cross-linked peptides.* Nat Commun. **10**(1), 3404.
2. Yang, B., Y.J. Wu, M. Zhu, S.B. Fan, J. Lin, et al. (2012) *Identification of cross-linked peptides from complex samples.* Nat Methods. **9**(9), 904-906.
3. O'Shea, J.P., M.F. Chou, S.A. Quader, J.K. Ryan, G.M. Church, et al. (2013) *pLogo: a probabilistic approach to visualizing sequence motifs.* Nat Methods. **10**(12), 1211-1212.
